# How to gain valuable insight from scarce data with Machine Learning: a post-hoc explanation tool to identify biases in biological images classification

**DOI:** 10.64898/2026.02.20.706981

**Authors:** Clémence Bolut, Anastasia Pacary, Laetitia Pieruccioni, Marielle Ousset, Jenny Paupert, Louis Casteilla, David Simoncini

**Affiliations:** University of Toulouse, Inserm, CNRS, EFS, ENVT, Restore Research Center, Toulouse, France; Université Toulouse Capitole, IRIT UMR5505, Toulouse, France; University of Toulouse, Inserm, CNRS, Infinity (Toulouse Institute for Infectious and Inflammatory Diseases), Toulouse, France

## Abstract

Machine learning (ML) models are effective at classifying images across various fields, including biology. However, their performance on biomedical images is often limited by the small size of available datasets that are constrained by the time-consuming and costly nature of experimental data collection. A review of the literature shows that many studies using biomedical images fail to follow ML best practices. This study focuses on regenerative medicine, which aims to promote tissue regeneration rather than scarring. To explore this process, we applied ML to a limited dataset of images of mice tissues, aiming to distinguish between regenerating and scarring samples. As expected binary classification failed to generalize to independent data. A novel SHAP-based analysis revealed that the overfitting models were based on spurious correlations including individual mice characteristics that aligned with the regeneration/scarring labels. The models appeared to be solving the binary classification task, but were in fact recognizing individuals. To investigate this behavior further, we examined the test set confusion matrix of a model trained to identify individual mice. We observed that, beyond individual recognition, individuals were grouped according to the time elapsed after injury (day 3 or 10) and the healing outcome (regeneration or scarring). We hypothesized that these groupings were based on relevant biological information captured by the model. To test this hypothesis, we successfully trained a model to classify images according to the time elapsed after injury (3 or 10 days), demonstrating that ML can extract relevant biological information when the task is aligned with what the data can actually support. Altogether, this study demonstrates that carefully examining explanations of a model is not only an effective way to unveil putative biases but also to extract relevant information from a limited dataset.

**Author summary:** Machine learning is increasingly used to analyze biomedical images, but in many experimental settings only small datasets are available, which can easily mislead powerful models. In this study, we looked at images from mice tissues, with the goal to distinguish healing by regeneration from healing by scarring. Although standard machine learning models appeared to perform well during training, they failed to generalize to new animals. By carefully analyzing model explanations, we found that the models were not learning biologically meaningful patterns of tissue repair, but instead were recognizing individual mice based on subtle image-specific signatures. Importantly, this same analysis revealed that the models did capture relevant biological information when the task was better aligned with the data, such as distinguishing early versus late stages of healing. Our results highlight how explanation methods can uncover hidden biases, prevent false conclusions, and help researchers extract meaningful biological insights even from limited and imperfect datasets.

## Introduction

Machine Learning (ML) provides tools to efficiently extract informative cues from large datasets in various disciplines including the biomedical field [1]. While Deep Learning (DL) and especially deep Convolutional Neural Networks (CNNs) enable to analyze biomedical images in order to automatically establish diagnostics or prognoses with high performances, these techniques can be limited in two ways. First, neural networks need large and diverse datasets to reach their full potential, a condition that is not easy to satisfy in biology where taking images involves manipulating living organisms, which is ethically questionable, time-consuming, difficult to standardize and expensive. Secondly, DL models trained on biomedical images can suffer from different biases with multi-factorial sources. Biases in ML have often been underestimated in all disciplines and especially in the biomedical field: while the number of DL papers increased exponentially since the 1990s and particularly between 2005 and 2020, several studies confirm that the impact of biases have long been minimized even for peer-reviewed manuscripts. For instance, a thorough examination of papers published in 2020 using ML techniques to detect and prognosticate Covid-19 from chest radiography or tomography images shows that no paper reaches the threshold of quality required for a realistic application in clinical practice due to data of poor quality, bad application of ML methodology, flaws in reproducibility or biases [2]. In another field, focusing on the validation process and data leakage in a corpus of papers about the diagnostic of Alzheimer’s disease using neural networks on magnetic resonance images of the brain demonstrated that half of these papers were biased due to data leakage [3].

Thanks to landmark discoveries in stem cell research, regenerative medicine makes it possible to regenerate injured tissues, while in adult mammals the repair process after an injury usually leads to scars and not to regenerated tissues. Though scarring is faster and more effective than regeneration for closing wounds, avoiding bleeding and preventing potential infections, it is always associated with at least a small alteration of the tissue function. Over time, these successive scars and tissue function alterations will participate to the emergence of chronic diseases and to the aging and decline of the individual [4]. A better characterization and early distinction of the two outcomes of healing, scars and regenerated tissues, could help better understand and monitor the tissue repair process in adult mammals, a mechanism that is currently not well comprehended. Often, the images used for such studies do not meet the usual conditions to efficiently train a ML model, being usually scarce and derived from a limited number of individuals.

Previously, we proposed an innovative preclinical model of healing that directs repair either towards scarring or regeneration, for a given tissue in adult mice with the same genetic background [5] [6] [7] [8]. Building on this established model, the present paper explores how a ML pipeline could provide an early prediction of tissue repair outcomes based on a limited number of minimal images of injured tissues. Particularly, we hypothesize that closely examining the models’ errors could help gain relevant insights from this imperfect data.

## Results

### The distinction between regenerating and scarring tissues fails to generalize

We first performed binary classification to distinguish images of regenerating from images of scarring tissues, using deep neural networks and XGBoost. The images were carefully placed in the training and test sets in order to avoid data leakage, as depicted in S1 Table.

Five-fold cross-validation was used for neural networks’ training. Trainings were repeated five times, with one fold being the validation set and the gathering of the four others the training set. Folds were built from the training set images of S1 Table, with all the 2D images from one 3D tile always kept together (but not necessarily all the tiles from one mosaic kept together). The test set was used to independently evaluate the trained neural networks.

In order to repeatedly train and test the XGBoost model on different configurations, five distinct datasets were built from the images of Table 2, each one made of a training and a test set. S1 Table corresponds to one of these datasets, the four others having similar numbers of images.

For each network architecture, Fig 1a shows the average loss and accuracy of the five models trained as part of the cross-validation process. The confusion matrices correspond to the evaluation on the test set of the models saved at the epoch with the lowest validation loss. Fig 1b presents the XGBoost averaged confusion matrices on the training and test sets.

**Figure 1.**
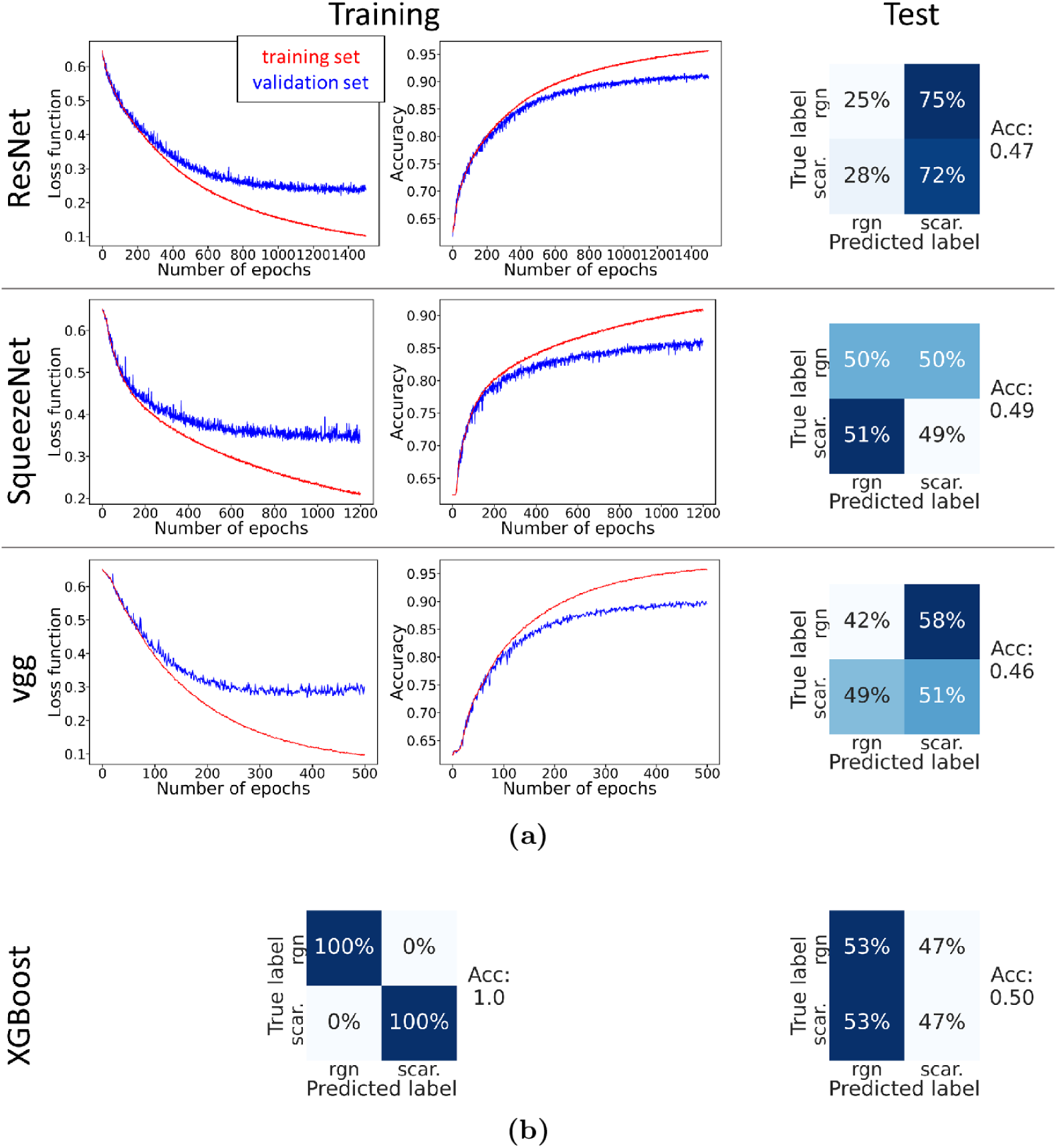
Regenerating/scarring classification with deep neural networks and XGBoost. **(a)** Three networks (ResNet, SqueezeNet and vgg) were used to classify the images between regenerating (“rgn”) and scarring (“scar.”). Values are averaged among the five cross-validation trainings. The first two columns show training and validation losses and accuracies. On the third column, confusion matrices evaluate the trained models on the test set, with corresponding accuracies (“Acc”); the first (resp. second) lines display the percentage of images predicted as one or the other class in relation to the total number of regenerating (resp. scarring) images. The results on the training and validation sets showed an apparent satisfying classification. But tests on an independent dataset demonstrated that the models did not generalize. **(b)** The XGBoost algorithm was trained for binary classification on features extracted from the images. Values are averaged among the five considered datasets. The left confusion matrix corresponds to the models’ predictions on the training sets, where the classification was perfectly learned. However, when tested on independent test sets, there was no generalization (right confusion matrix) and an accuracy similar to random guesses.

For the three considered neural networks architectures, both training and validation losses decreased. Over the successive epochs training and validation losses diverged, prompting training termination to prevent overfitting. Similarly, both training and validation accuracies increased over time, eventually plateauing near 1, which indicates improved models’ predictions on both sets. However, this apparent successful classification was contradicted by the confusion matrices on the test set which revealed that none of the architectures correctly classified the images and had overfitted. Here the data was averaged among the five trainings of cross-validation, so in order to obtain a refined analysis we computed the accuracies and F_1_-scores of all trained models on the training and test sets (S1 Fig, with also XGBoost data). The training and validation accuracies showed little variance, indicating consistent results across the five folds; they ranged from 0.75 to 0.97, denoting significantly better than random (0.5) predictions on these sets. However, a significant drop was observed between training and test accuracies, with all test accuracies settling at 0.5. These equivalent-to-random predictions on the test set indicated overfitting: the apparent learning seen on the training and validation sets was not able to generalize well on independent data.

Similarly to what was observed with neural networks, the XGBoost model seemed to properly learn to classify between regenerating and scarring images in the first place, but failed to generalize. Indeed, the training confusion matrix demonstrates that XGBoost was effectively trained on this set where it achieved perfect classification. However, when tested on an independent test set, the confusion matrix shows that the model seemed to randomly guess each image’s class, with an average accuracy of 0.50.

Overall, these results reveal that the binary classification was not learned correctly whatever ML model was considered, suggesting the presence of a bias that prevents generalization to an independent test set.

### Efficient individuals classification can be achieved

Given the limited number of individuals and the intrinsic biological heterogeneity between them, we hypothesized that if the models were able to discern mice, then it could represent a strong bias towards the distinction between regenerating and scarring tissues. To investigate this hypothesis, we trained the models to distinguish between the individual mice.

Because mice classification’s goal was to evaluate whether the tiles of one mosaic (corresponding to one mouse) could be differentiated from the tiles of another mosaic (another mouse), the test set needed to contain some images of all mice, contrary to binary classification. 90% of the tiles of each mouse were randomly chosen for the training set and the remaining ones were placed in the test set, leading to the distribution shown in S2 Table. Its training and test sets were used to train neural networks with five-fold cross-validation, similarly as for binary classification. Also, the XGBoost model was trained on five dataset configurations from a subset of these images. Indeed, to better balance the classes we excluded from the 22 available individuals the five mice with the smallest numbers of tiles and we removed some tiles of the mouse with the biggest number of images, resulting in a dataset containing the images of 17 mice.

Fig 2a shows the losses (first column), accuracies (second column) and confusion matrices on the test set (third column) averaged among the five models of cross-validation. Fig 2b displays the XGBoost averaged confusion matrices on the training and test sets.

**Figure 2.**
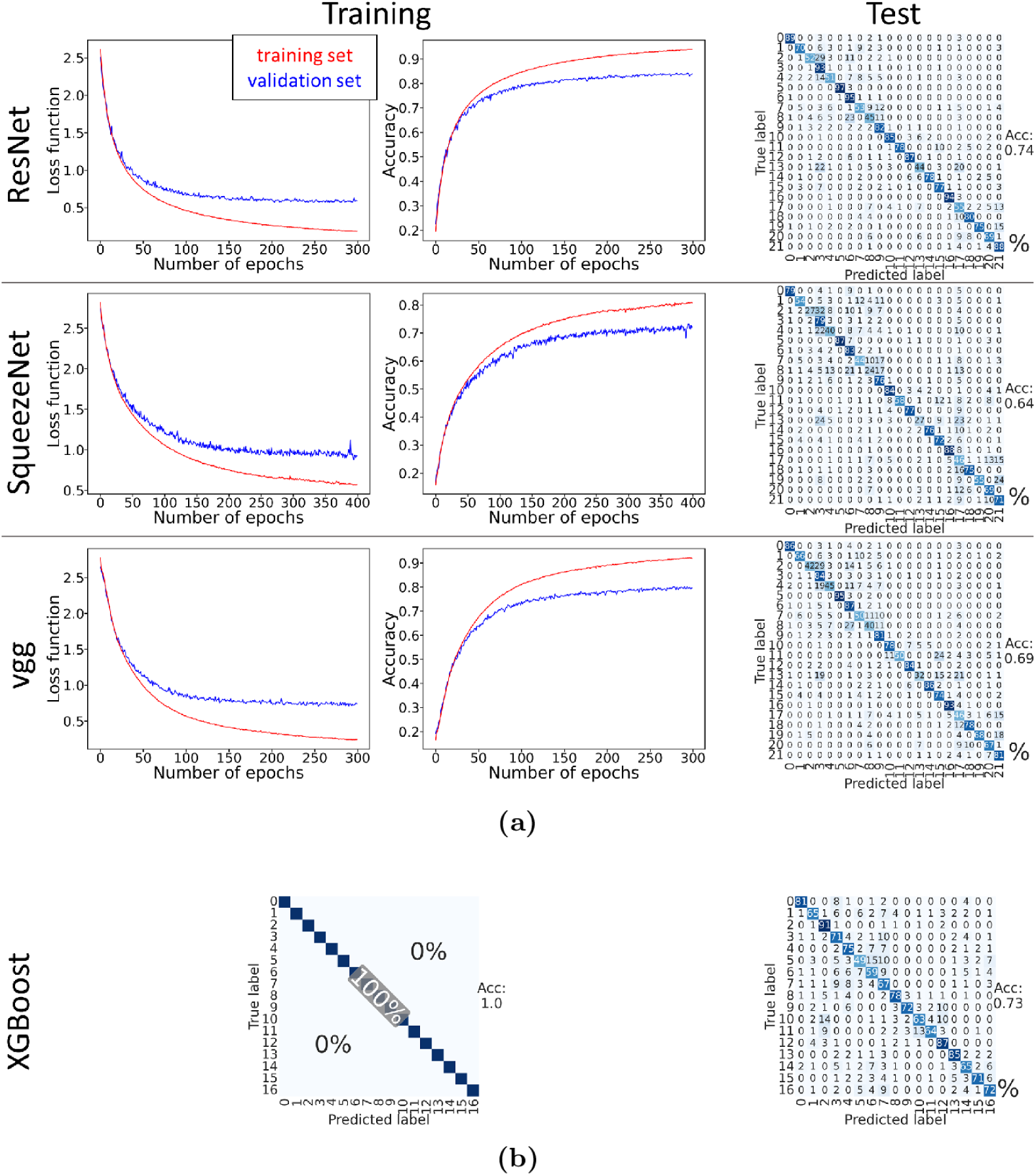
Mice classification with deep neural networks and XGBoost. **(a)** Three networks were used to classify the images between the different mice. The averaged losses, accuracies and confusion matrices on the test set are displayed. On the confusion matrices, the value at position (*i, j*) is the percentage of images of mouse *i* predicted as mouse *j*, in relation to the total number of images of mouse *i*. The three networks learned properly: the losses decreased, the accuracies increased towards 1 during training and the mice were still recognized from one another on images of the test set never seen by the networks before. **(b)** The XGBoost model was trained for mice classification on features extracted from the images. Values are averaged among the five considered datasets. The left confusion matrix shows the performances of the model on the training sets, where it provided perfect predictions. The right confusion matrix corresponds to the classification on the test sets and shows that the model did properly generalize to independent data.

For the three neural networks architectures considered, the training and validation losses decreased during training and the training and validation accuracies increased towards 1, suggesting that the networks classification power improved during training. The accuracies of the five models trained as part of the cross-validation process displayed in S2 Fig showed little variance on the training set, meaning that the results among the five folds were homogeneous. At the end of training, training accuracies between 0.75 and 0.89 were obtained for ResNet, SqueezeNet and vgg, which was significantly higher than a random guess of probability 1*/*22 ≃ 0.05. These performances still held on the test set. Therefore, ResNet, SqueezeNet and vgg were able to efficiently distinguish to which mouse a given image belonged.

The results of the XGBoost training (see S2 Fig for the values of all trained models) were similar to the ones obtained with neural networks. The training set confusion matrix shows that the model was able to learn to classify between the different mice. The test set matrix confirmed that it could generalize to new images, with an averaged accuracy of 0.73 significantly higher than the random guess of probability 1*/*17 ≃ 0.06.

These results show that distinguishing between mice is a rather straightforward task that can be successfully accomplished using either a neural network or XGBoost.

### SHAP analysis highlights strong similarities between binary and mice classifications

We then examined whether the XGBoost models trained on binary classification relied or not on the same key features as the models trained on mice classification, in order to better evaluate the similarities between the two tasks. To do so, the SHAP (for *SHapley Additive exPlanations*) explainability tool [9] [10] was used to analyze one of the five XGBoost models trained for binary classification and one of the five XGBoost models trained for mice classification.

For binary classification, a single SHAP plot (top of Fig 3) was sufficient to get the ranking of the most important features for this task. But for the mice recognition task we generated one SHAP plot per mouse (S3 Fig). To obtain a single measure of feature importance for the entire mice classification task, we defined two metrics. These metrics summarized the information from the seventeen SHAP plots into a single feature ranking. The two metrics were the average position of the features in the SHAP plots and the number of occurrences of the features in the SHAP top 20s. The features that were ranked high by the two metrics were the most important features for recognizing mice according to SHAP (see bottom of Fig 3 and S4 Fig).

**Figure 3.**
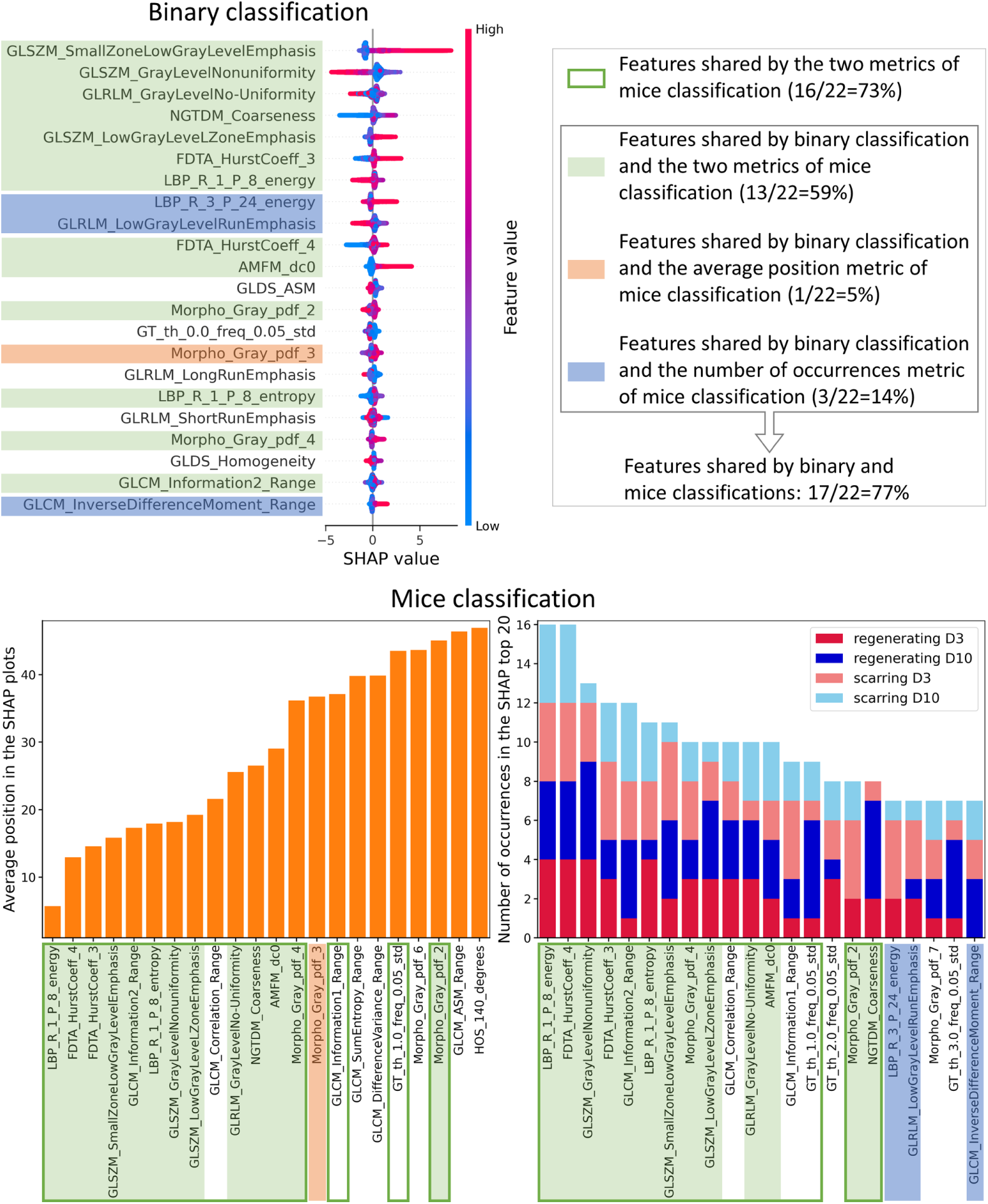
Regenerating/scarring classification’s crucial features are similar to the ones of mice classification. The 22 most important features of binary and mice classifications are shown. **Above**: SHAP feature ranking for one of the XGBoost models trained for binary classification. **Below**: two metrics were used to provide an overall ranking according to SHAP of the most important features for one of the XGBoost models trained for mice classification. The most important features for mice classification were the 16 features shared by the two metrics (here bordered by a green line). **Comparison**: there were 77% of common features in the highest ranked features of binary and mice classifications. The common features are highlighted in color: green features are shared by the binary classification and the two metrics of the mice classification (59% of the most important features), orange (resp. blue) ones by the binary classification and the average position (resp. number of occurrences) metric of mice classification (5% and 14% of the most important features).

This analysis with SHAP shows that XGBoost models trained for binary and mice classifications were based on the same crucial features to make their predictions. The two classifications had 17 shared features in total, representing 77% of all of the 22 features considered, or 13 shared features (59%) if we consider the more stringent condition to be in the binary classification ranking and in the two rankings of mice classification.

We further compared the XGBoost models trained for binary and mice classifications by examining their dependence plots for the most important features shared by the two tasks. As S5 Fig demonstrates that the dependence plots were highly consistent with the results described earlier (non-generalizing binary classification and proper mice classification), dependence plots can be regarded as particularly relevant to investigate our models’ behavior. Fig 4 presents, for feature GLSZM_SmallZoneLow GrayLevelEmphasis, the dependence plot of binary classification to the left and overlaid views of the dependence plots of mice classification to the right. An analogous comparison for two other crucial features was carried out in S6 Fig.

**Figure 4.**
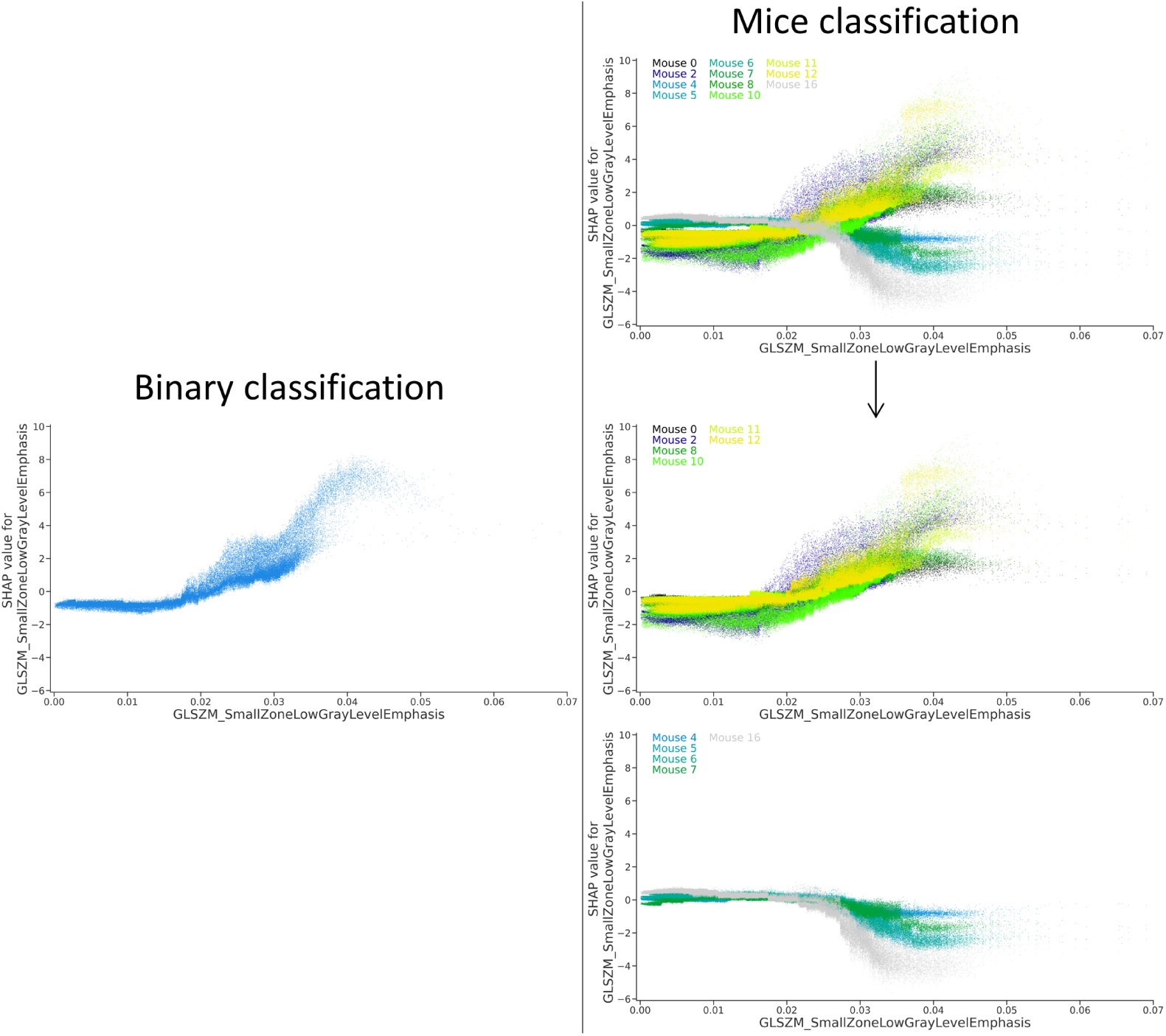
Comparison of profiles between regenerating class dependence plot and mice classification dependence plots of the regenerating mice. Dependence plots of feature GLSZM_SmallZoneLowGrayLevelEmphasis for the XGBoost models trained for both classifications. **Left:** binary classification, SHAP values for the prediction of “regenerating”. **Right:** for mice classification, the global overlaid view (first line) shows the superposition of the dependence plots for the prediction of all the mice. The dependence plots of all regenerating (resp. scarring) mice were similar, their layering is displayed on the second (resp. third) line. **Comparison:** the dependence plot of the XGBoost model trained for binary classification resembled the superposition of the dependence plots of all the regenerating mice.

On the global overlaid view for mice recognition, the dependence plots of each mouse could be distinguished from the others: each branch was separable and had its specific shape. This was consistent with the fact that the considered feature was one of the most important for mice classification and thus enabled to particularly well differentiate mice from one another. Moreover, the overlaid view revealed two distinct curve patterns, allowing to group mice into two corresponding subgroups (lines two and three). These subgroups corresponded to different repair outcomes, with the first group comprised solely of regenerating mice and the second one including only scarring mice. So the regenerating mice had similarly looking dependence plots, and the scarring mice as well. Therefore, feature GLSZM_SmallZoneLowGrayLevelEmphasis, which accurately discriminated between mice, also provided clues about the repair class of the images. Furthermore, the binary classification dependence plot for the prediction of class “regenerating” closely resembled the mice classification dependence plots of the regenerating mice subgroup. This suggests that the way the “regenerating” class was predicted by the model in the context of binary classification was similar to the way the regenerating mice were recognized in the context of mice classification. Overall, the similarities observed between the dependence plots of the two classification tasks indicate similarities in the way the two tasks were carried out by XGBoost.

### Error analysis reveals data subgroups on which unbiased learning is performed

To further improve our understanding of binary classification’s failure, we thoroughly examined the errors made by XGBoost when classifying mice, and more specifically the mistakes distribution regarding the four biological subgroups highlighted in red in Fig 5. On the left of this figure are displayed the XGBoost confusion matrices on the tests sets of the five dataset configurations (their average is displayed in Fig 2b), and on the right the averaged percentage of images that were predicted in a wrong subgroup.

**Figure 5.**
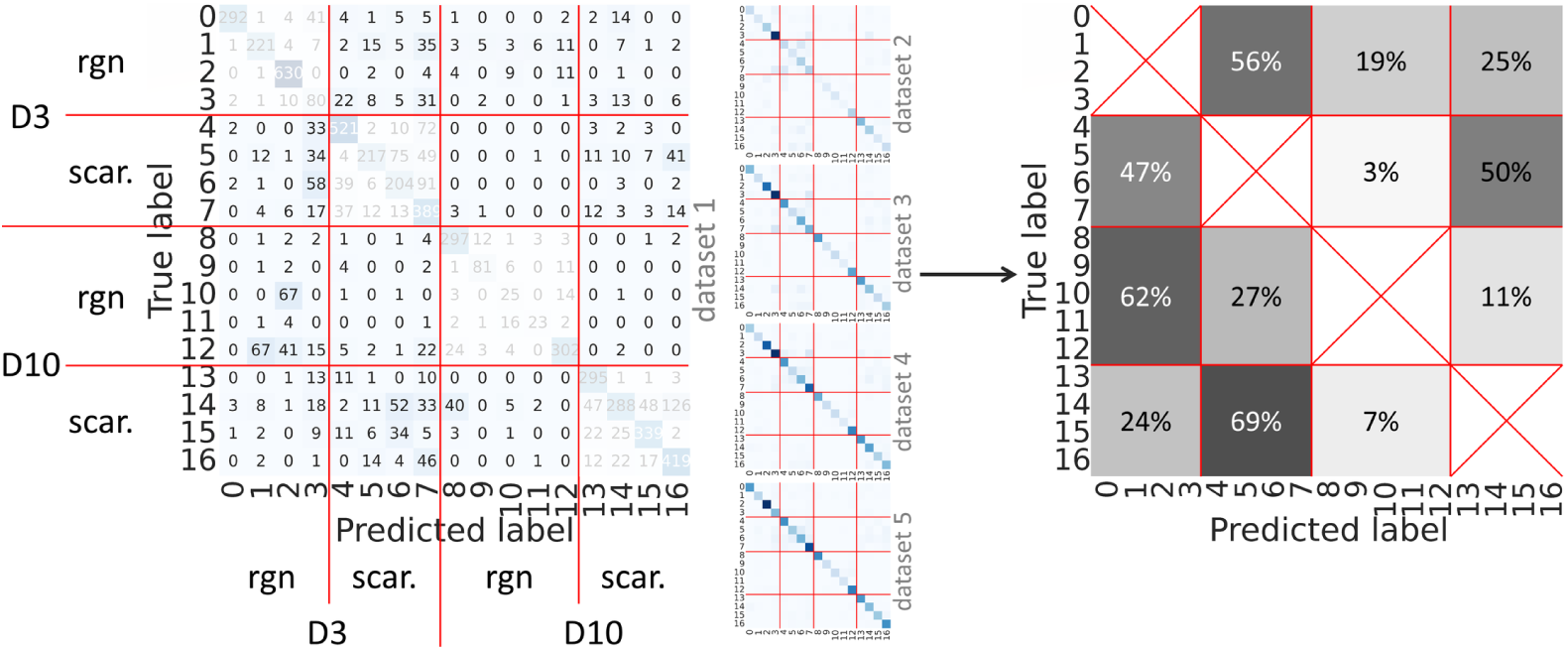
Repartition of errors on mice classification. **Left:** Confusion matrices on the test sets of XGBoost trained on the five dataset configurations for mice classification. The four mice subgroups depending on the repair class and on the time of tissue sampling are indicated with red lines: D3 regenerating (“rgn”) mice (four individuals), D3 scarring (“scar.”) mice (four individuals), D10 regenerating mice (five individuals) and D10 scarring mice (four individuals). **Right**: Prediction errors for each subgroup, averaged among the five dataset configurations. On the D3 regenerating line, the three cells indicate the percentage of images wrongly predicted in another subgroup than D3 regenerating: among all the errors made for D3 regenerating images, 56% were wrongly predicted for D3 scarring, 19% for D10 regenerating and 25% for D10 scarring. The same calculation was made for the three other subgroups.

Prediction mistakes were not completely random and showed distinct patterns across the four subgroups. For instance, mispredictions of D10 mice were made most often towards D3 mice of the same class: D10 regenerating mice images were mostly misclassified for D3 regenerating mice (62%) and D10 scarring for D3 scarring mice (69%). So the errors distribution of mice classification was not uniform but in fact linked with the four biological subgroups.

Guided by this mistakes analysis, we investigated in detail whether SqueezeNet and XGBoost were able to achieve unbiased binary classification on combinations of the four subgroups (see Table 1 for their ability to generalize to independent test sets). Both models were trained on the six possible image combinations of the subgroups, with the images of S1 Table. Depending on the chosen combination, the models were trained to distinguish between regenerating and scarring images or between D3 and D10 images.

**Table 1.**
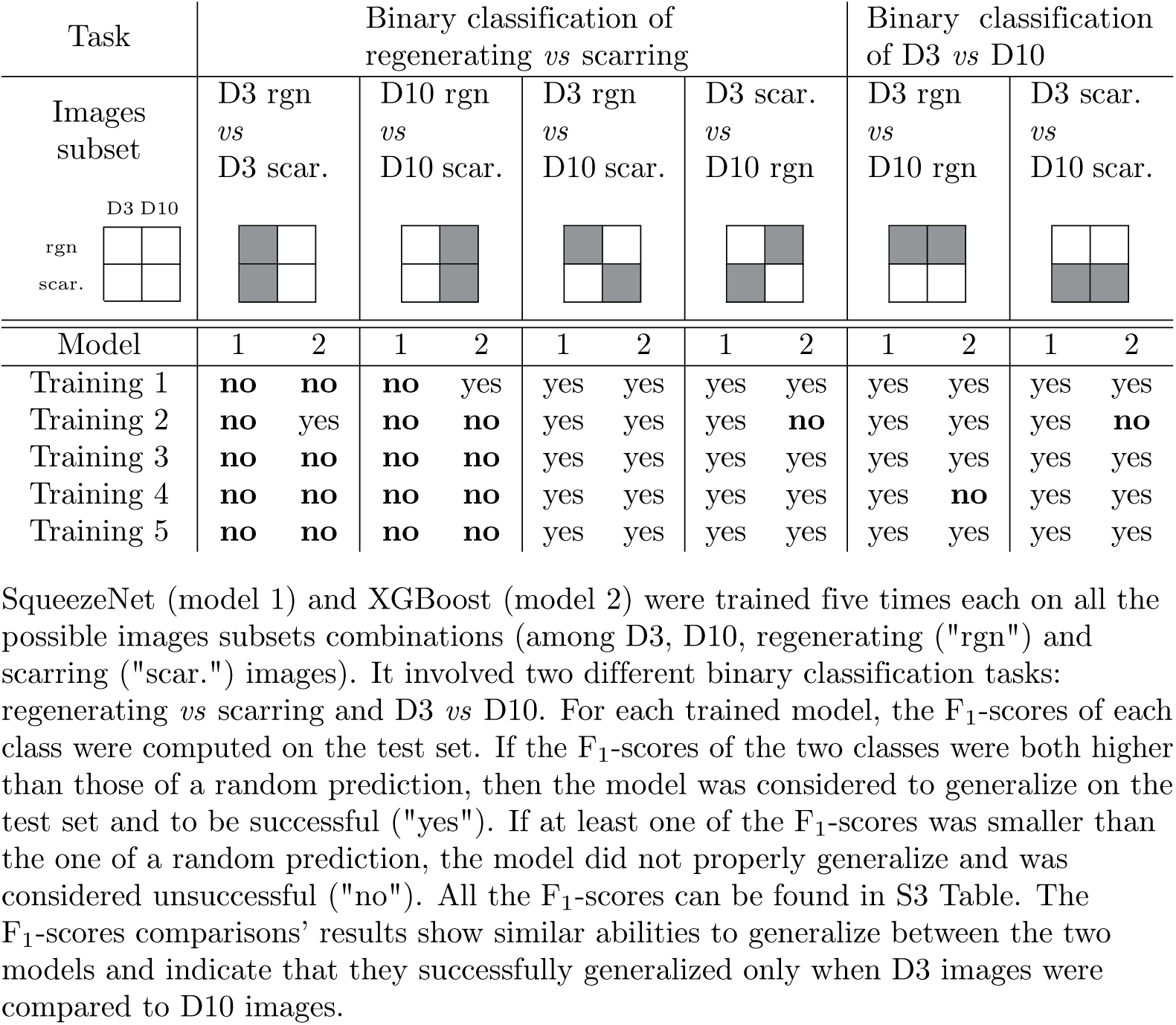
Classification with a deep neural network and XGBoost only generalizes on the subsets mixing D3 and D10 images.

The F_1_-score was used to assess the ability of a trained model to generalize to an independent test set. For each model, its F_1_-score on the test set for each class (either regenerating and scarring or D3 and D10) was compared with the F_1_-score of a random classifier (see S3 Table for all values). A model was considered successful (*i.e.* capable of generalizing to an independent dataset) only if the F_1_-scores of the two classes were both higher than those of a random guess.

The results are consistent within the five trainings and the two models, with successful trainings for the classifications “D3 regenerating *vs* D10 scarring”, “D3 scarring *vs* D10 regenerating”, “D3 regenerating *vs* D10 regenerating” and “D3 scarring *vs* D10 scarring”. In the first two cases, the groups to distinguish had both distinct tissue repair types (regenerating/scarring) and different tissue sampling times (D3/D10), and it was therefore unclear what drove the distinction. However, the inability to discern D3 (or D10) regenerating from scarring images suggested that the success in “D3 regenerating *vs* D10 scarring” and “D3 scarring *vs* D10 regenerating” was driven by tissue sampling times rather than repair types. This was confirmed by the successful binary classification of D3 *vs* D10, with the distinction of D3 and D10 regenerating images (and similarly for scarring).

Overall, the analysis on the subsets demonstrates that although the neural network SqueezeNet and the XGBoost model failed to differentiate regenerating from scarring images, which was our original goal, they both successfully succeeded in learning some insightful biological features of the images: the distinction between D3 and D10 images.

## Discussion

Biomedical research often requires to collect data through experiments involving animals. However, the number of animals used is typically limited due to ethical considerations as well as financial and logistic constraints. As a result, the available data is often scarce and fails to adequately represent the biological diversity relevant to the task. DL models are valuable because they can capture complex non-linear dependencies, helping to identify key mechanisms that even trained human experts might overlook, thereby accelerating research and advancing science. This advance and the widespread availability of foundation models have further increased both the use and misuse of powerful ML techniques in biomedical research, with misuses mainly arising from biases in the original and limited datasets. The classic misuse in the field comes from data leakage when training and test sets are not properly constructed. But models may also pick up on random, non-generalizable patterns that happen to correlate with the target labels in the training data. These spurious correlations, with potentially several features involved across different data splits, can lead to deceptively good performance during evaluation, while in fact preventing the model from generalizing to new data. Identifying and mitigating such effects is critical to ensure robust conclusions.

Our study focused on data from the field of regenerative medicine, which aims at guiding tissue repair towards regeneration rather than scarring. To investigate this process, we used a limited number of minimal images to train ML models at predicting regenerating and scarring tissues in mice. Our images were available in few amounts, they stemmed from a restricted number of individuals and they were not very diverse (the tiles of one mosaic looked similar between themselves).

Initially, when considering only the training set, the models appeared to efficiently learn to classify the images. However, testing the models (neural networks and XGBoost) on a dataset entirely independent from the training data revealed that the classification between regenerating and scarring tissues was unable to generalize. Remarkably, although this requirement of test set independence has been extensively discussed in the literature [11] [12] [13] [14], it is often not properly addressed in many studies across various application domains [15] [3] [16] [17].

It was shown previously with the experiment “Name That Dataset” that when using several public datasets for object recognition tasks, each dataset tends to display its own identifiable and unique signature [18] [19]. In order to explore our data inter-individual variability (heterogeneity between mice), we classified individual mice. Mice classification was key to understanding biases and identifying conditions under which the tissue repair binary classification (regererating/scarring) could succeed. Indeed, although both neural networks and XGBoost overfitted and failed to satisfyingly perform binary classification, they were able to predict the identity of individual mice using the same limited dataset. This suggests that, rather than learning features relevant to the tissue repair outcomes, the models were biased toward recognizing individual mice. This hypothesis is consistent with the way the datasets were constructed: for binary classification different mice were used to provide the images of the training set and of the test set, while for mice classification images from the same mice were split between the training and *validation* sets in each cross-validation configuration.

To explore the connection between both classification tasks, we proposed a novel use of the post-hoc explanation tool SHAP. SHAP was employed to study whether the most important features used for classification were shared between the two tasks and, through comparison of dependence plots, to investigate potential similarities in the way features influenced predictions. Our analysis showed that the same key features were used in both the tissue repair binary and the mice identity classifications and that the dependence plots were highly similar, supporting our hypothesis that the tissue repair classification was biased by individual mice recognition. It appears that the similarities among tiles from the same mouse were easier for the models to learn than the actual repair status of the tissues, revealing hidden biases.

A thorough examination of the mice identity classification revealed a structure that led us to define an alternative, biologically relevant task that could be learned from the same limited dataset. Carefully inspecting the model’s errors of classification through its test confusion matrix demonstrated that they were not randomly distributed, but rather followed patterns linked to the four biological subgroups (thus when the model wrongly predicted D10 mice, it mostly mistook them for D3 mice of the same repair class). This error analysis suggested that, despite the bias in the tissue repair classification, the models did capture some biologically meaningful characteristics of the images.

Based on this insight, we tried new classifications and demonstrated that, using the same dataset, D3 images could be successfully distinguished from D10 images. This conclusion aligns with recent transcriptomic comparisons [20] (unpublished) indicating that differences between days in the repair process (*e.g.* D3 and D10) are more pronounced than those between the regenerating and scarring outcomes.

In conclusion, our analysis shows that spurious correlations can be identified by thoroughly examining two tasks: the learning of the original target and the identification of individuals. The tasks can be ingeniously analyzed with local explanation models and the impact of the most predictive features can be compared. This approach enables to isolate features that have strong, validated predictive power for individual recognition but that also, by chance, split the training set relatively well according to the original target labels. Besides helping to eliminate spurious correlations, our analysis of the individual recognition task also brought to light a learnable biological phenomenon, even though the dataset used was limited and biased. One promising outcome of this analysis is the development of ML pipelines that can seamlessly discard features leading to spurious correlations. It would enable to denoise the data, improve learning, but also extract relevant biological information even based on small datasets from a limited number of individuals.

## Methods

### In vivo models

All experiments were undertaken on seven-week-old C57BL/6 OlaHSD male mice coming from Envigo Laboratories. Animals were reared in conventional conditions at the Faculty of Pharmacy in Toulouse: mice were grouped at three to five per cage, they were bred in a regulated environment (21°C, twelve-hour light/obscurity cycles) with an unlimited water access and a standard diet. Cervical dislocation was used to kill the mice. The ARRIVE guidelines of the European Community Council and the European Community Guidelines (2010/63/UE) were respected. All animal experiments were authorized by the Ministry of Higher Education, Research and Innovation and by the French ethics committee.

At day zero, all mice underwent unilateral resection of their subcutaneous adipose tissue (AT, see S7 Fig). After isoflurane 2.5% was inhaled by the animals for anesthesia, their right AT was accessed through a single abdominal incision. No surgical procedure was performed on the left AT that was used for internal control.

For some mice, 35% of the right AT was excised, between the lymph node and groin (normal size resection). For other mice, 15% of the right AT was excised (small size resection); to precisely locate the resection area, the same points of reference as the normal size resection we used, pliers were put between these points and the cut was made on either side of it. In both cases, the skin was closed with three suture points.

From days zero to three after AT resection, mice treatment consisted in daily subcutaneous injections of 50 *µ*L of naloxone methiodide (NaL-M, 17 mg/kg, N129, Sigma Aldrich, for normal size resections) or NaCl (for small size resections).

Three or ten days after resection, mice were killed to retrieve the part of the tissue that was cut beforehand. Samples of mice AT three or ten days after resection were thereby obtained, where each sample was labeled as “regenerating” or “scarring”.

### Imaging procedure

The AT samples were fixed (PFA 4%, 24h), incubated for 1.5 hour in PBS (0.2% triton) at room temperature, stained with DRAQ5 dye (Thermofisher 62251 used at 5mM, a dye that intercalates into DNA) for two hours in the dark and finally mounted between slide and slip cover.

Images were acquired with a confocal biphoton microscope (confocal microscope LSM880 from Zeiss with a coherent biphoton laser and an objective lens LCI “Pan Apochromat” 10X/0.45 or “LD C-Apochromat” 40x/1.1) with two channels to observe the cells’ nuclei (stained with DRAQ5, *λ*_excitation_ = 633 nm, *λ*_emission_ = 650 nm) and the collagen fibers (second-harmonic generation, *λ*_excitation_ = 800 nm, *λ*_reflected_ = 400 nm). All acquisitions were made with a magnification of 40x.

The acquired images were 3D images (*tiles*) made of several 2D slices separated by 0.5 *µ*m or 5 *µ*m; each 3D tile was made of 30 to 366 2D images depending on the tissue’s depth. Each image was a square of side length 212 *µ*m (512 × 512 pixels). Each pixel’s value was encoded on 16 bits. Several 3D images were taken for each sample, forming a mosaic of tiles with no overlap between them on each tissue. Each mouse provided multiple images, consisting of mosaics of varying sizes (ranging from 1 to 257 tiles per mouse) and tiles of different depths. As a result, the total number of images per mouse varied significantly, with some producing 90 images and others up to 7394. Altogether, samples coming from 28 different mice were imaged (15 regenerating and 13 scarring mice), corresponding to 48788 2D images with two channels.

The methods about the *in vivo* models and imaging procedure are similar to the ones of [21].

### Data preprocessing

Firstly, the software Fiji (https://fiji.sc/) was used to automatically convert the images’ Zeiss format *czi* to the Python-handable *tiff*. Because the confocal microscope was tuned to acquire automatic mosaics of the AT samples, some images were totally black or contained only uninformative noise. Two thresholds were defined: the minimal value of a pixel for it to be considered bright enough (20 for fibers, 9 for nuclei) and the minimal proportion of bright pixels in an image for it to be deemed informative (5 × 10*^−^*^5^ for fibers, 2 × 10*^−^*^4^ for nuclei). All the images bellow the second threshold were automatically deleted and 31% of the initially acquired images were retained for the next steps.

The deep-learning-specific preprocessing that was used consisted in cropping and normalizing the images. As most of the images displayed a darker area in the 100 pixel column on the right of the image, all the images were cropped from 512 × 512 pixels to a 400 × 400 pixels square centered to the left. As the pixels were encoded on 16 bits and thus varied between 0 and 2^16^, all the pixels’ values were normalized to [0; 1] by dividing them by 2^16^. This cropping and normalization procedure was used in the exact same way for the two channels (fibers and nuclei).

The images were then split in two sets: training and test. 2D images from the same 3D tile were all put in the same set. For binary classification, images from the same mouse were placed either in the training or in the test set.

### Neural networks training

Three different neural networks architectures were used: ResNet18, SqueezeNet and vgg11. The first layers were changed to allow for two-channel images.

The data augmentation pipeline consisted in random cropping (from the 400 × 400 pixels images to 250 × 250 pixels), flipping (with 0.5 probability), and changes of brightness and contrast (by choosing a random brightness and contrast value in [0; 2]).

Five-fold cross-validation was implemented on the training dataset that was used for training and validation.

Depending on the number of classification classes, either cross entropy loss or binary cross entropy loss was used. Models were trained using Stochastic Gradient Descent (SGD) as optimizer. The batch size was always 16 and there was a 10*^−^*^5^ weight decay. The learning rate was *lr* = 10*^−^*^3^, except for ResNet trained for binary classification (*lr* = 10*^−^*^4^).

Models were trained using the computing centers Calmip and Occidata (Toulouse, France).

### XGBoost training

Features were extracted from the images using the PyFeats library [22], generating for each image 2682 features across 24 categories (textural, morphological, histogram and multi-scale). The Boruta feature selection algorithm [23] was used to reduce this to 911 features for binary classification and 1057 for mice classification. The XGBoost algorithm was then trained on these features using optimal parameters determined via gridsearch, with performances monitored with confusion matrices on training and test sets.

### SHAP analysis

The trained XGBoost models were analyzed with the TreeSHAP explainability method, which orders features by importance and provides quantitative links between features’ values and predictions [9].

For binary classification, one SHAP plot was sufficient to indicate the most important features for predicting the two outcomes. For mice classification, we obtained as many SHAP plots are there are mice, we thus summarized all this information into a general list of the most important features.

The influence of the features’ behavior on the prediction was examined through dependence plots. For the prediction of a certain class, the dependence plot of a given feature indicates which SHAP values are assigned to the images depending on the values of this feature. For binary classification, one dependence plot was enough to reveal the links between a given feature and the model’s predictions. But for mice classification, for each feature as many dependence plots as there are mice could be drawn. In order to get a global view of a feature’s influence on the model, the dependence plots of all the mice were put on top of each other with different colors. Therefore, a dependence plot for binary classification shows each image once, while for mice classification on the global overlaid view each image is represented as may times as there are mice. However, in order to avoid overloading the mice classification dependence plots, only mice with sufficient influence on the considered feature (being in the top 20 most important features for this mouse’s prediction) were drawn on the overlaid view. The obtained dependence plots can be analyzed further by representing the images’ labels with different colors.

### Complete pipeline and obtained dataset

As previously described, the model of AT resection is particularly relevant to investigate repair processes in adult mammals [5] [6] [7] [8] [24] [25]. In this model, scarring and regenerative healings of adult mice are respectively induced with a small size resection and with an early and transient treatment with an antagonist opioid. When regeneration occurs, the complete process is achieved two months after resection. In this model, the issue of tissue repair was subject to some heterogeneity estimated at around 70% (30% of the mice treated with naloxone injections will not regenerate) [5]. We investigated whether ML applied to a limited number of minimal images of tissues three (D3) or ten (D10) days after resection could provide an early prediction of the final outcome of tissue repair (Fig 6).

**Figure 6.**
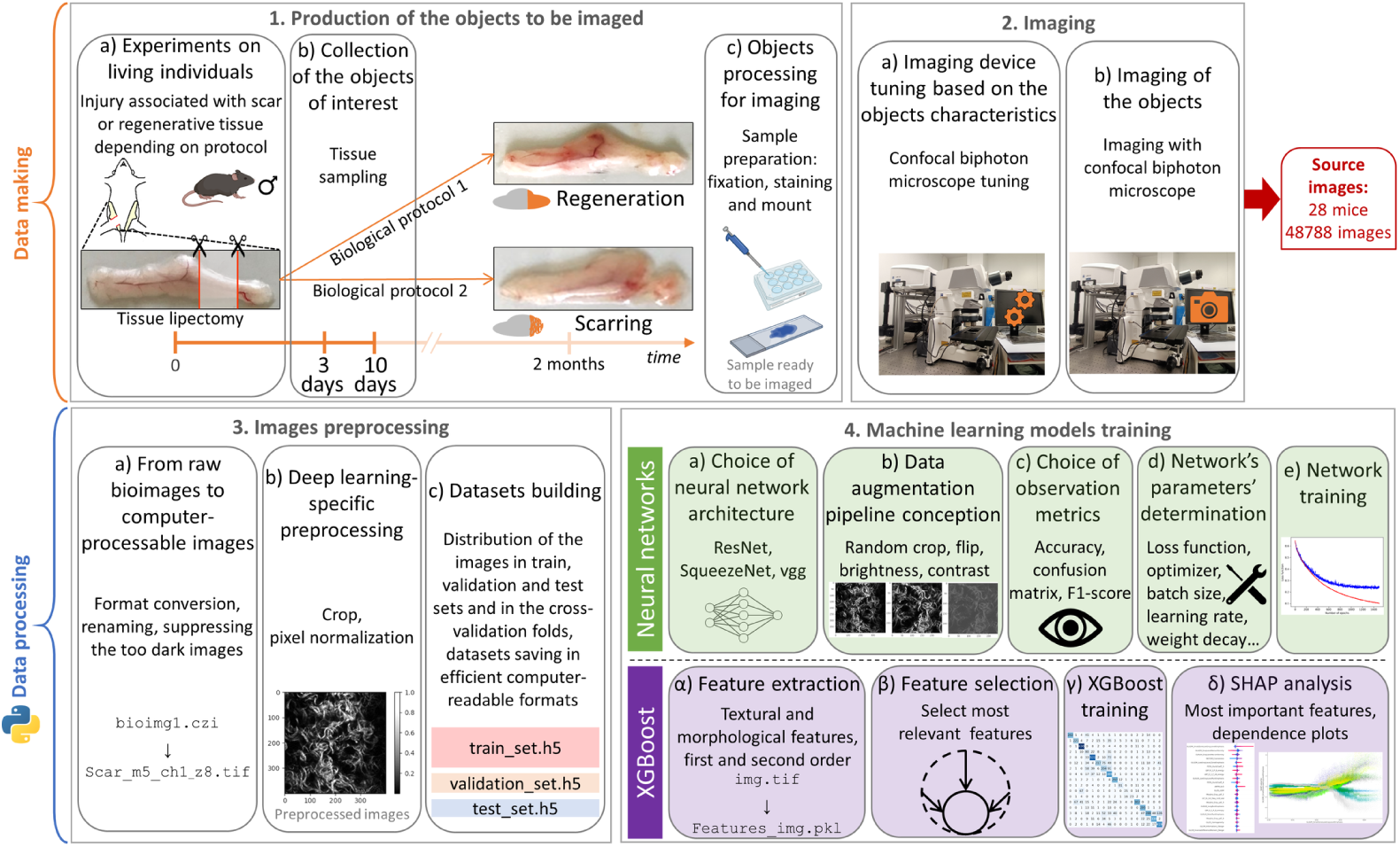
End-to-end ML pipeline. After surgery, mice were treated to induce regenerating or scarring healing. Three or ten days after surgery, the tissues were collected, prepared for imaging and imaged. A confocal biphoton microscope was used to generate the source images that constituted the main dataset. This raw data was then preprocessed and distributed in the datasets used for training, validation and testing. After that, two ML techniques were used: neural networks and XGBoost followed by an analysis with SHAP.

Images stemmed from the tissue samples of 28 mice divided into four subgroups. The characteristics of all the 48788 acquired images are depicted in Table 2.

**Table 2.** Characteristics of all acquired images.

|  |  | Number of mice | Number of 2D images (with two channels) | Average number of 2D images per mouse |
| --- | --- | --- | --- | --- |
| D3 | regenerating | 6 | 10863 | 1811 |
|  | scarring | 6 | 12681 | 2114 |
| D10 | regenerating | 9 | 8162 | 907 |
|  | scarring | 7 | 17082 | 2440 |
| subtotal regenerating |  | 15 | 19025 | 1268 |
| subtotal scarring |  | 13 | 29763 | 2289 |
| **total** |  | **28** | **48788** | 1742 |

For each mouse, several 3D images (tiles) were acquired, each tile being made of several 2D images with two channels (fibers and nuclei).

Because the 2D images arising from a 3D tile might be quite similar and because the tiles of one mosaic (*i.e.* of one tissue of one mouse) might resemble each other, we carefully built datasets to avoid data leakage by keeping in the same set (training or test set) all the 2D images stemming from the same 3D tile. Two classification tasks were performed on these images: binary and mice classifications. The building of the training and test sets, which depended on the classification task, is described in the Results section.

## Ethics Statement

All animal experiments were carried out in compliance with European Community Guidelines (2010/63/UE) and approved by the institutional ethics committee and by the Ministry of National Education, Higher Education and Research (protocol reference: #40880-2023010612201954 v6).

## Acknowledgments

We thank Mathieu Vigneau and Corinne Lorenzo from the Center for Expertise and Technological Resources (CERT platform), and Thomas Serre and his team from Brown University, Providence (RI, USA) for insightful comments and discussions.

This work was founded by the Imperial College London - CNRS PhD Joint Programme 2020 (Grant 80PRIME). This study was partially supported through the grant EUR CARe n°ANR-18-EURE-0003 in the framework of the “Programme des Investissements d’Avenir”. This work was granted access to the HPC resources of the CALMIP supercomputing center (CALcul en MIdi Pyrénées, Toulouse, France) under the allocations 2016-P16043 and 2016-P24042, and to the computational platform Occidata of the Toulouse Institute for Computer Science Research (IRIT).

## Supporting information

**S1 Table.**
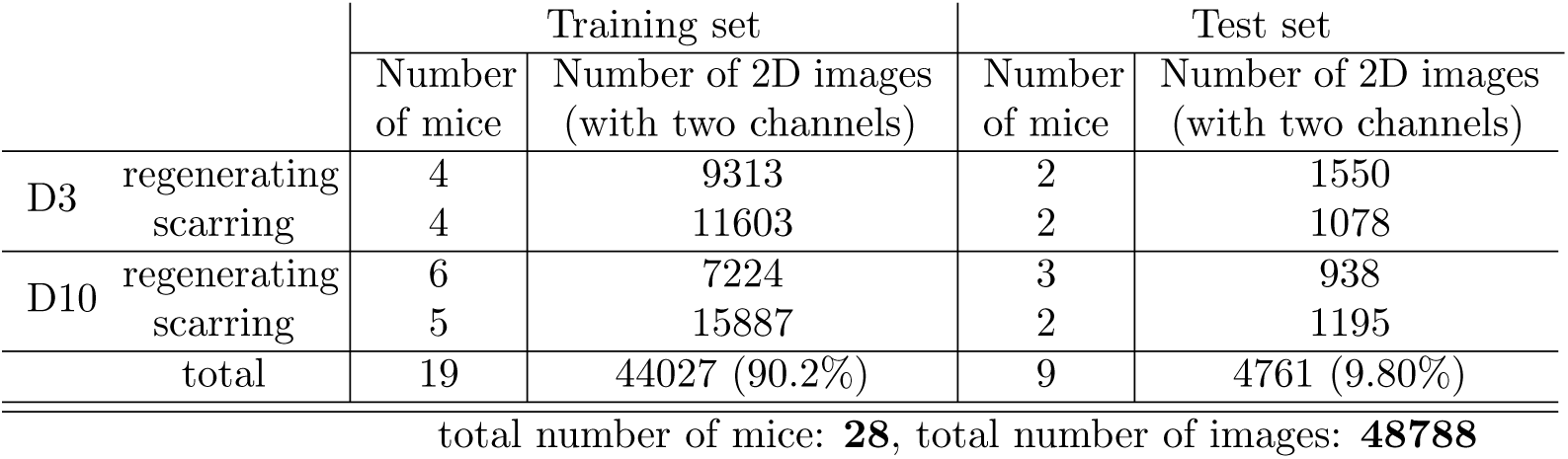
Training and test sets for binary classification. Mice from the training and from the test sets were not the same individuals. All the images acquired on one tissue sample (*i.e.* on one mouse) were put either in the training set or in the test set, so that all the tiles of a specific mouse could be found only in one set. This is to prevent data leakage because the tiles of a given mouse might be similar with each other as they were acquired in a row with the same microscopy parameters.

**S1 Fig.**
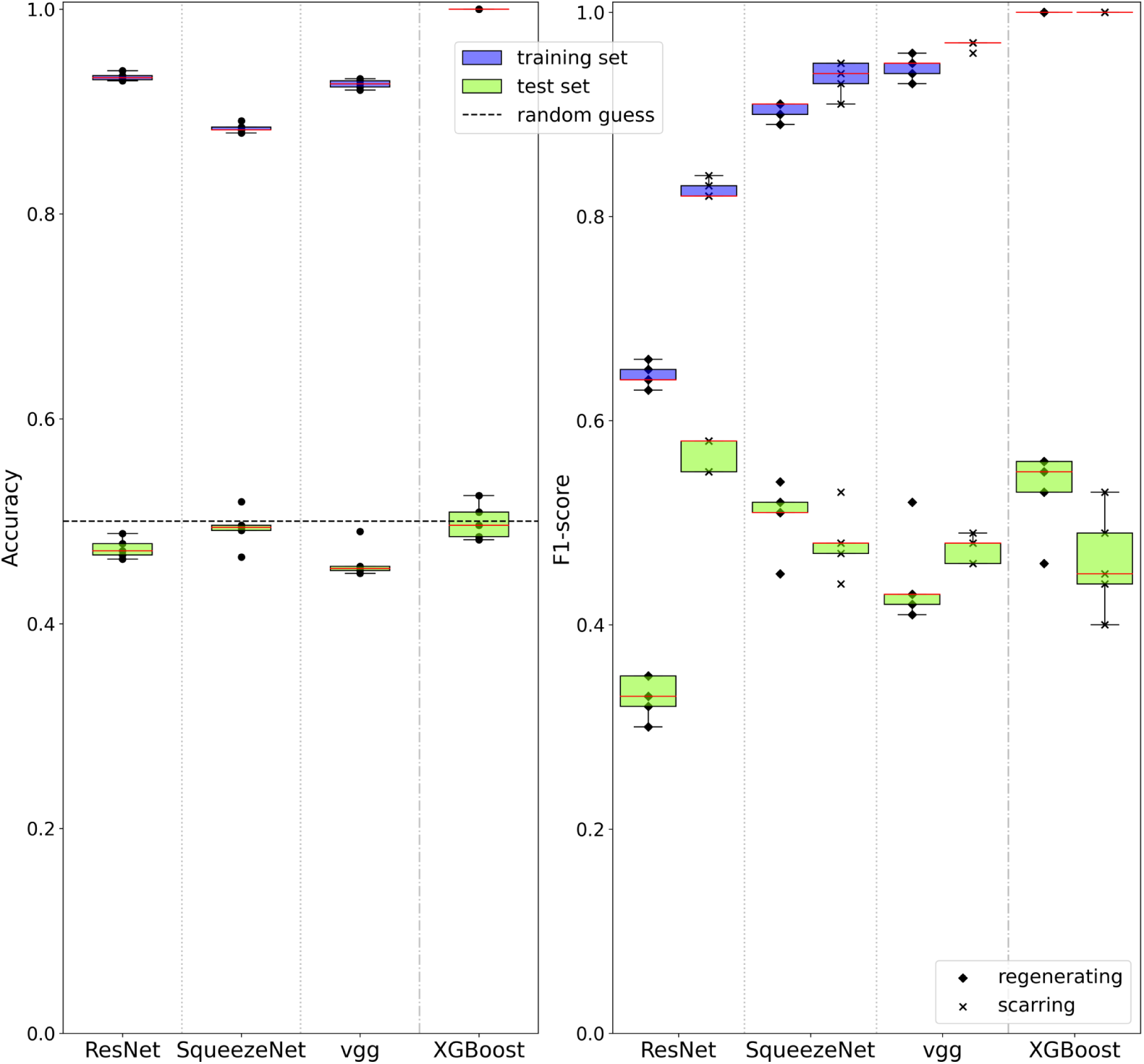
Evaluation of the models (three neural networks and XGBoost) trained for binary classification. Each dot corresponds to one of the five neural network models trained as part of the five-fold cross-validation process, or to one of the five dataset configurations considered for XGBoost. Fig 1 shows results averaged among these multiple trainings. **Left**: the accuracies of all models are displayed on the training and test sets. For the three neural networks architectures, there was a drop in accuracy between the training and test sets with test accuracies close to 0.5 corresponding to random guesses. This indicates the incapacity of learnings made on the training set to generalize to an independent test set. Similarly, the trained XGBoost models achieved perfect classification on all the five training sets and a random accuracy close to 0.5 on the test sets. **Right**: F_1_-scores for the two classes (“regenerating” and “scarring”) are displayed for the same models and sets as to the left. The performances of the four models on the training data appeared to be similar when considering only the accuracies. However, a closer look at the F_1_-scores revealed that for SqueezeNet, vgg and XGBoost the F_1_-scores for “regenerating” were close to the ones for “scarring”, while for ResNet they differed more. This difference in prediction between the two classes for ResNet, not observed for the other models, indicates a degraded performance of the former architecture.

**S2 Table.**
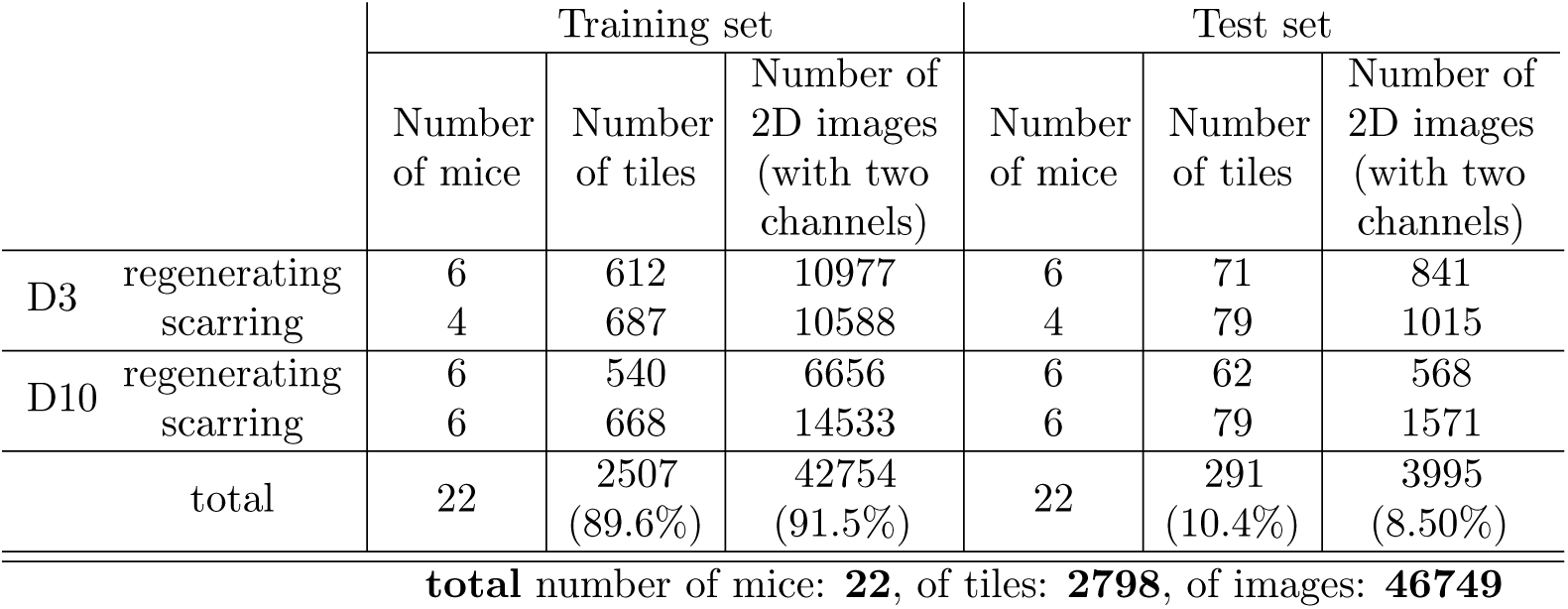
Training and test sets for mice classification. Among the 28 mice of Table 2, the 6 individuals which did not have complete mosaics acquired on them (for them, only less than ten non-adjacent tiles were acquired) were irrelevant for mice classification and therefore excluded from the datasets. Mice from the training and from the test sets were the same individuals. For each tissue sample (*i.e.* each mouse), some of its 3D tiles were put in the training set and the other tiles in the test set.

|  |  | Training set |  |  | Test set |  |  |
| --- | --- | --- | --- | --- | --- | --- | --- |
|  |  | Number of mice | Number of tiles | Number of 2D images (with two channels) | Number of mice | Number of tiles | Number of 2D images (with two channels) |
| D3 | regenerating | 6 | 612 | 10977 | 6 | 71 | 841 |
|  | scarring | 4 | 687 | 10588 | 4 | 79 | 1015 |
| D10 | regenerating | 6 | 540 | 6656 | 6 | 62 | 568 |
|  | scarring | 6 | 668 | 14533 | 6 | 79 | 1571 |
| total |  | 22 | 2507<br>(89.6%) | 42754<br>(91.5%) | 22 | 291<br>(10.4%) | 3995<br>(8.50%) |

**total** number of mice: **22**, of tiles: **2798**, of images: **46749**

**S2 Fig.**
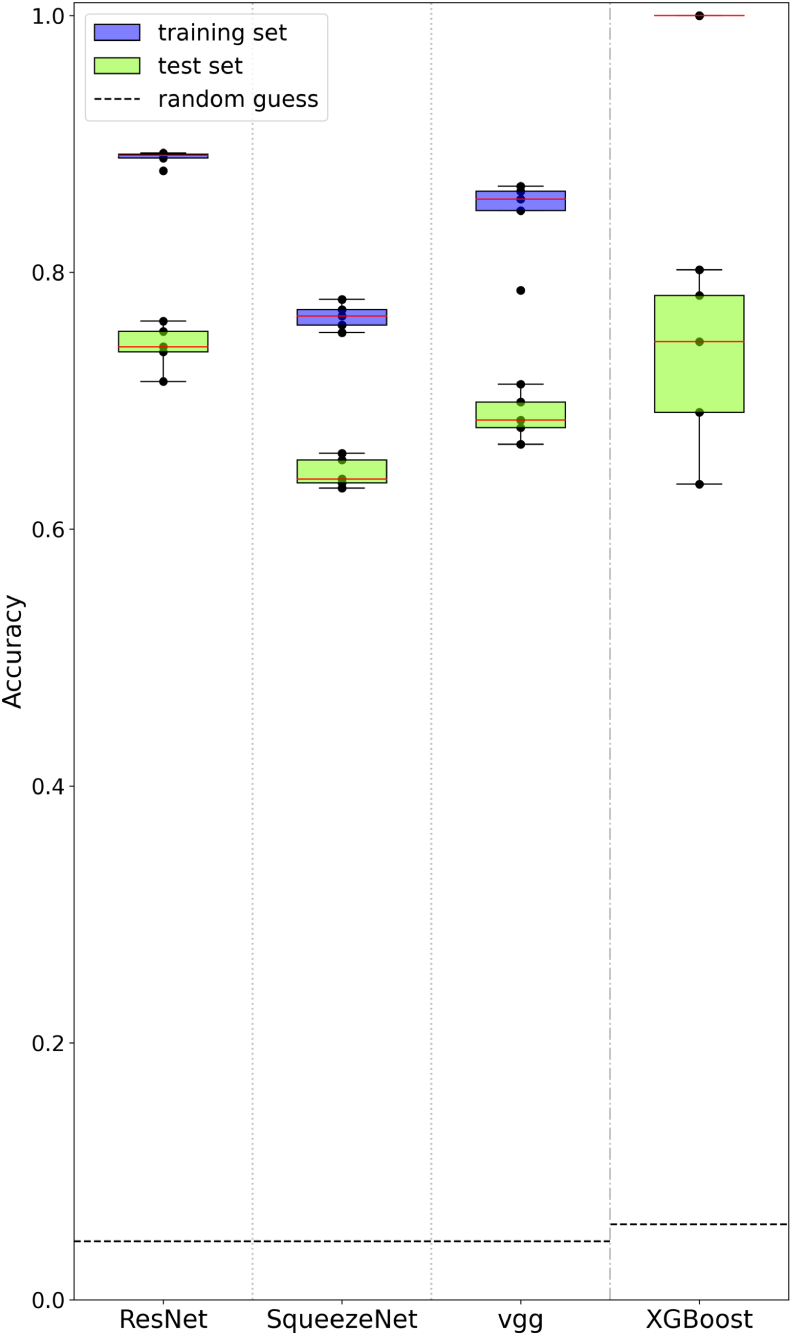
Evaluation of the models (three neural networks and XGBoost) trained for mice classification. Each dot corresponds to one of the five neural network models trained as part of the five-fold cross-validation process, or to one of the five dataset configurations considered for XGBoost. Fig 2 shows results averaged among the multiple trainings. The accuracies of all models are displayed on the training and test sets. For the three neural networks, the accuracies were close between the training and test sets. All the test sets displayed an accuracy significantly higher than random guesses (1*/*22 ≃ 0.05). The trained XGBoost models achieved perfect classification on all the five training sets and an accuracy significantly higher than random guesses (1*/*17 ≃ 0.06) on the test sets. The learning was successful for all the four models.

**S3 Fig.**
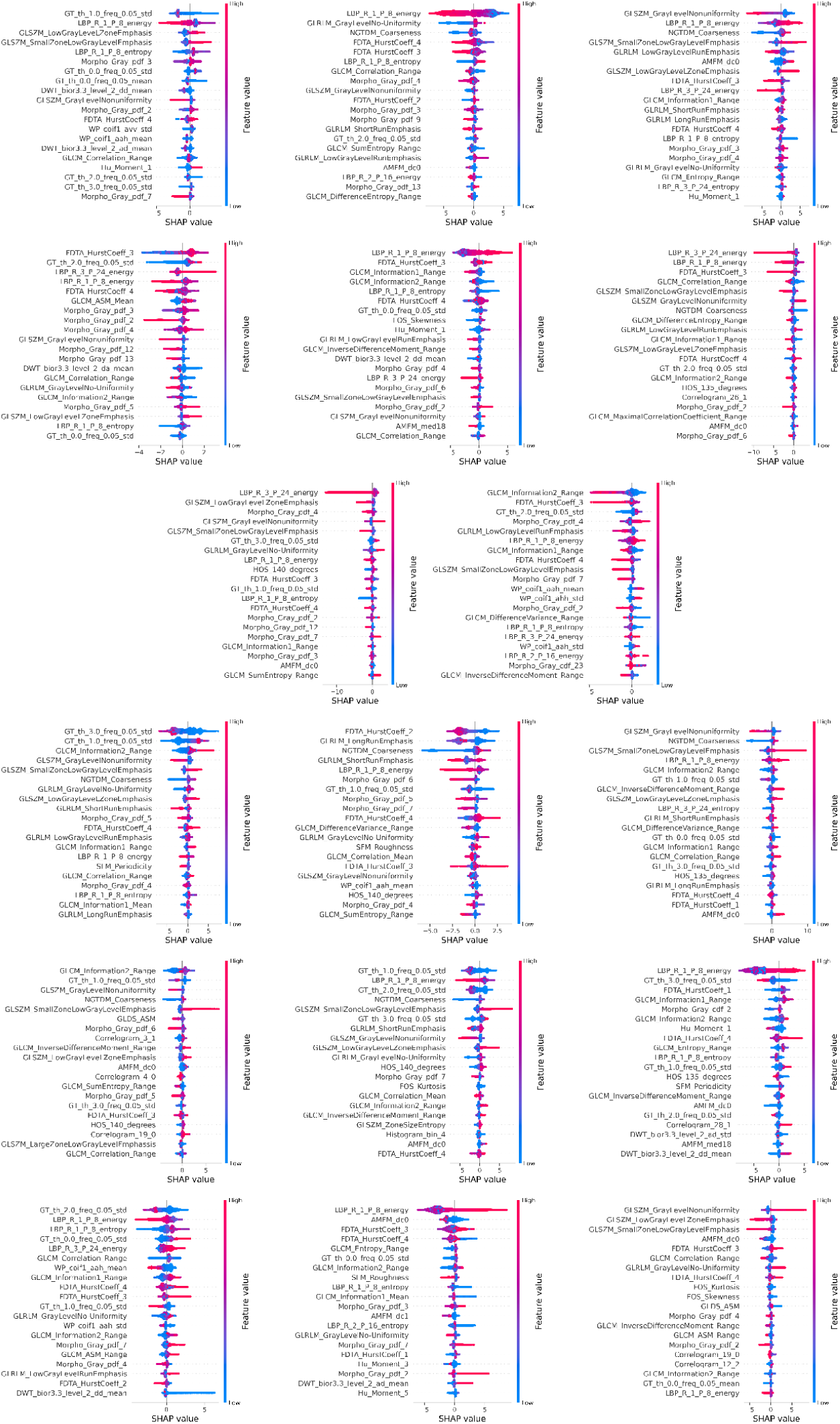
SHAP plots to explain one of the XGBoost models trained for mice classification. Each SHAP plot shows the most important features for the classification of a given mouse. The first three lines correspond to the D3 mice (four regenerating and four scarring mice) and the last three lines to the D10 ones (five regenerating and four scarring mice).

**S4 Fig.**
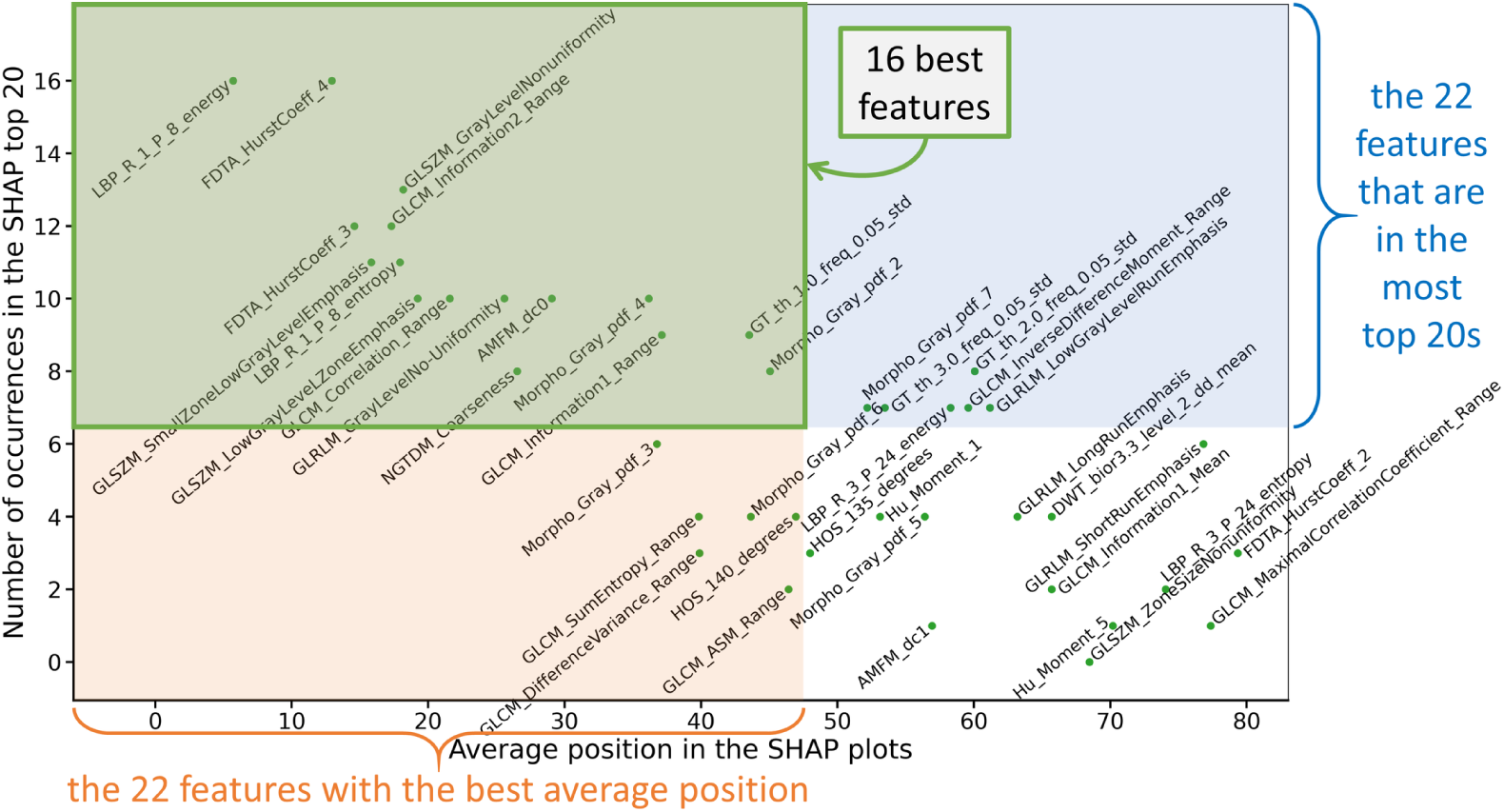
The 16 most important features for mice classification according to SHAP. Analysis of the SHAP plots of the 17 mice shown in S3 Fig with two metrics on the features: their average position in the SHAP plots and their number of occurrences in the top 20s. The 16 most important features for mice classification are highlighted in green (see also the bottom of Fig 3).

**S5 Fig.**
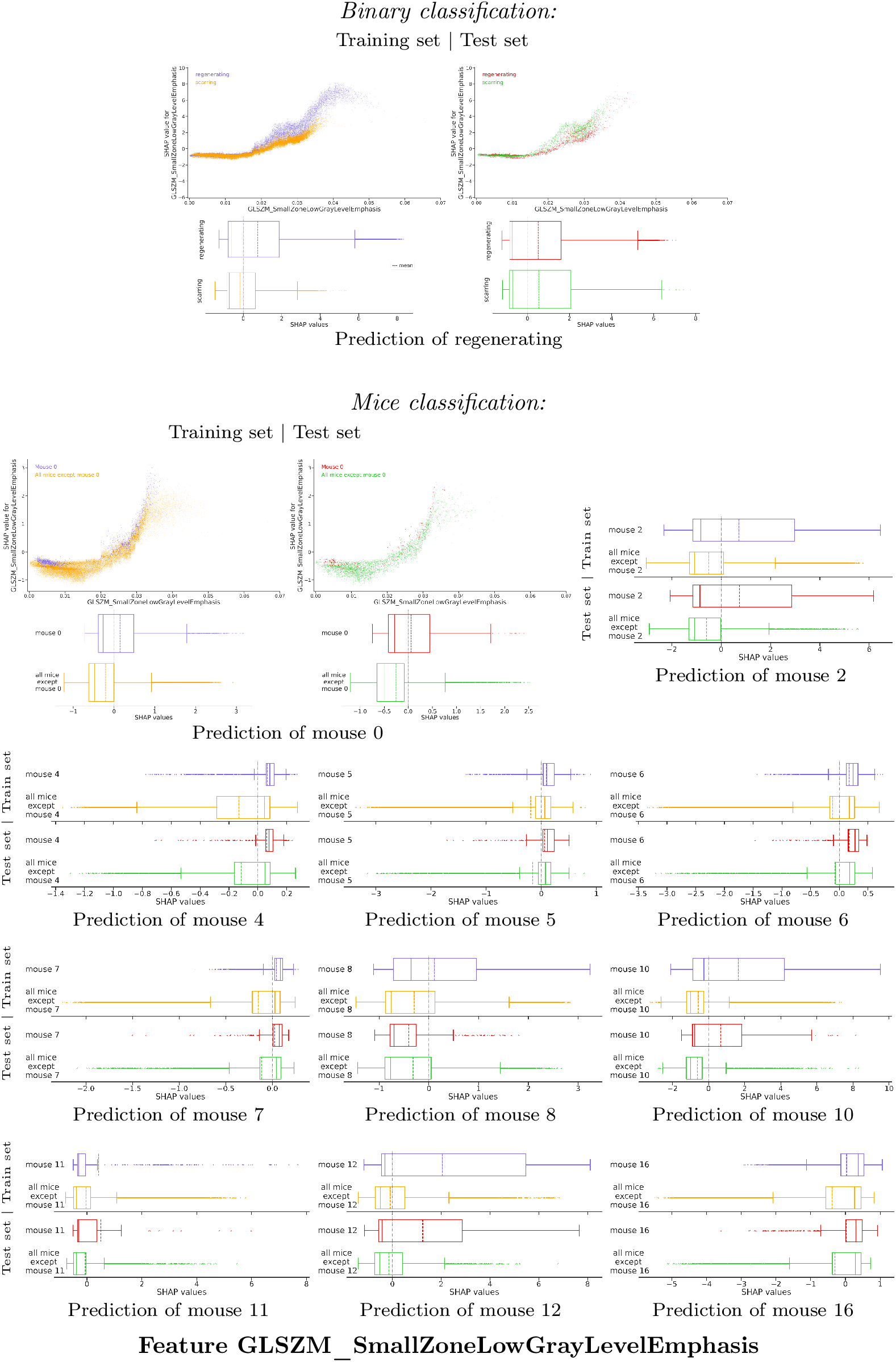
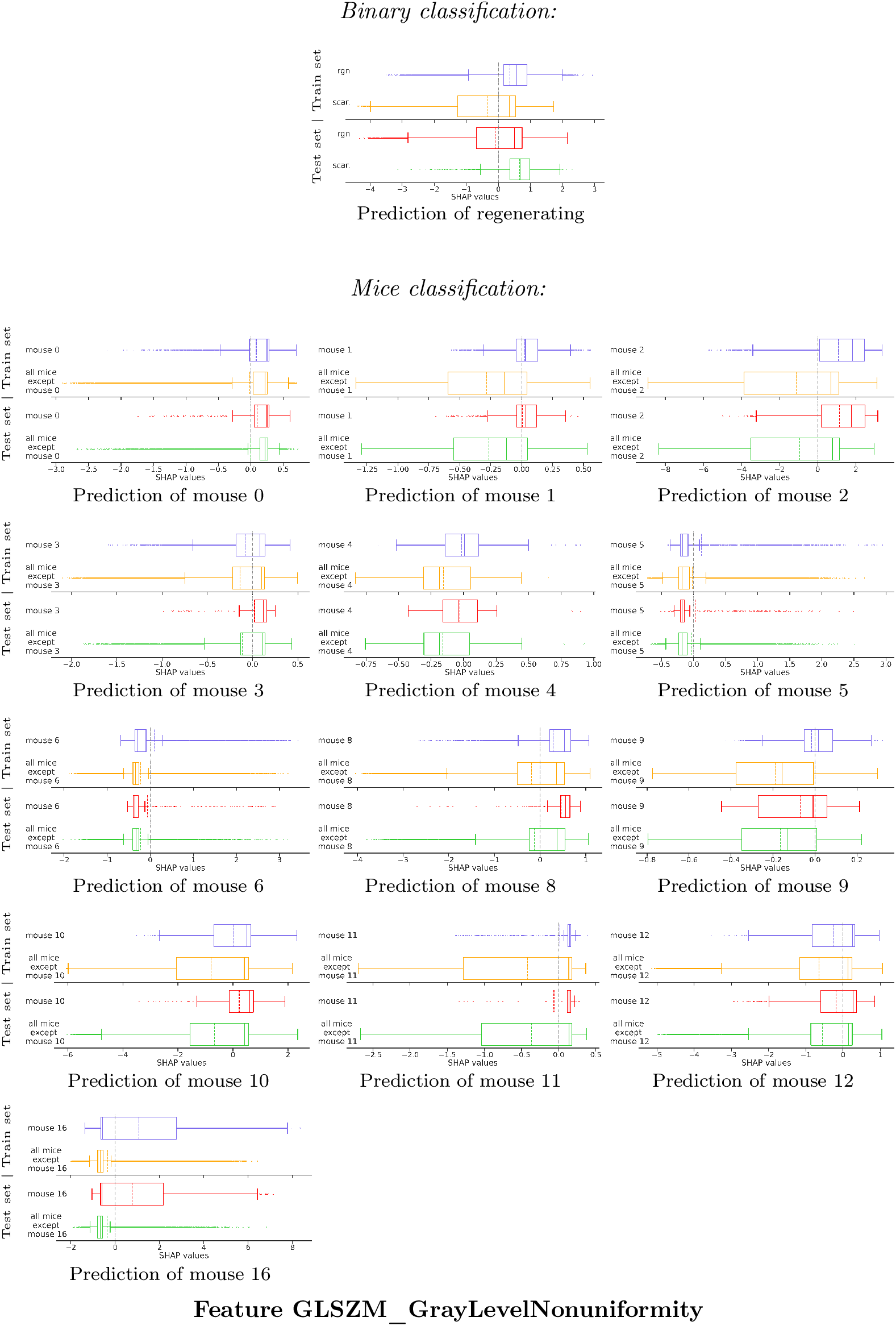
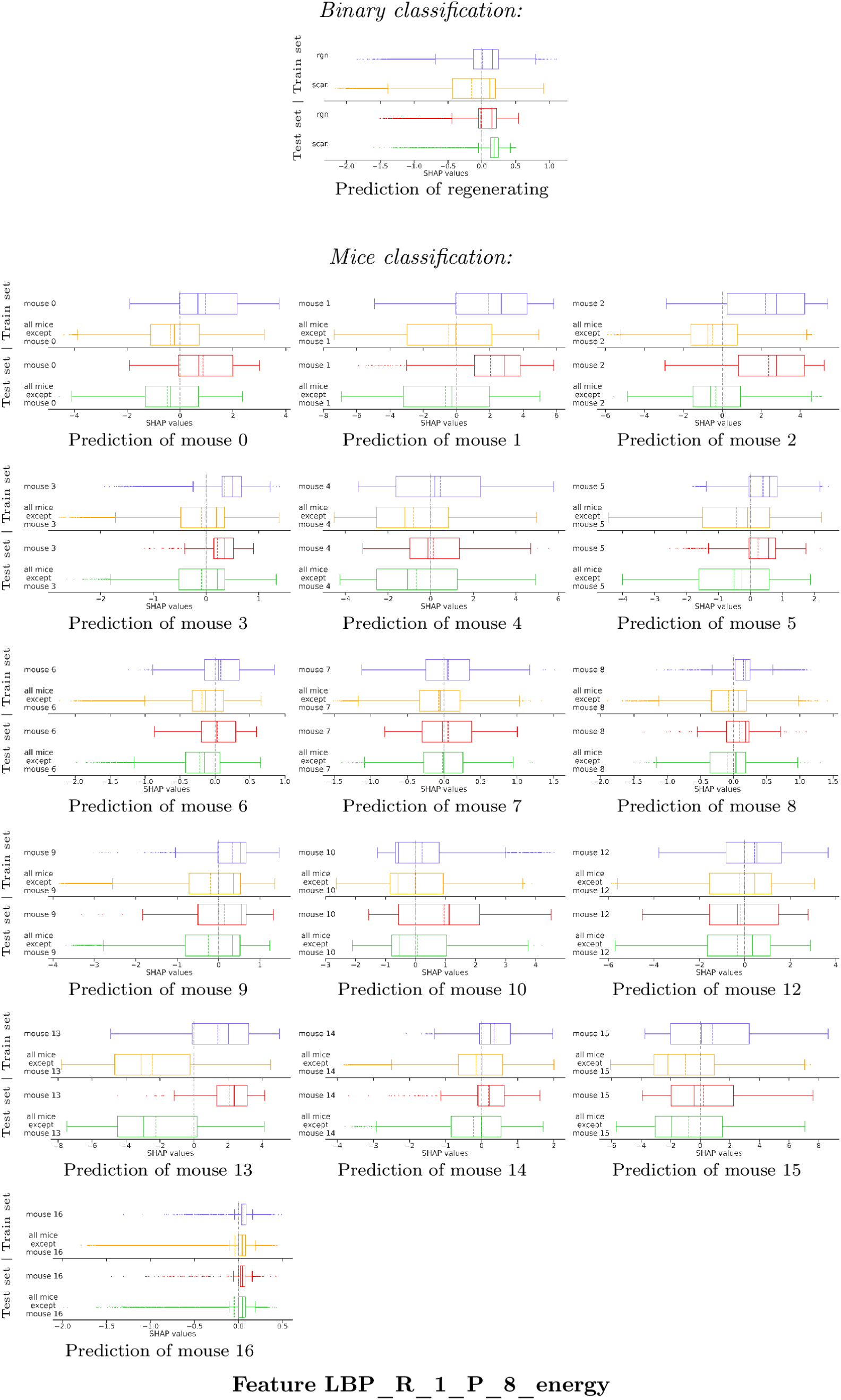
The dependence plots of the most important features display consistent non-generalizing binary and generalizing mice classifications. For the XGBoost models trained for binary and mice classifications, analysis of the SHAP values for features GLSZM_SmallZoneLowGrayLevelEmphasis (most important feature for binary classification as indicated in Fig 3), GLSZM_GrayLevelNonuniformity (second most important feature for binary classification) and LBP_R_1_P_8_energy (most important feature for mice classification) using dependence plots (only shown twice for concision) and corresponding box plots (with means displayed with dotted lines) that quantitatively measure the distribution of the classes. Training set (lines one and two of each subfigure) and test set (lines three and four) images were drawn separately. For the prediction of a given class, two colors were used: one for this class and another for all the other classes. For all features and for binary classification, regenerating (“rgn”) images had overall higher SHAP values than scarring (“scar.”) ones on the training set, and similar values on the test set. For mice classification on both sets for the prediction of mouse *m*, the images of mouse *m* had higher SHAP values than the images of all other mice. This is consistent with the non-generalizing binary classification and generalizing mice classification shown in Fig 1b and Fig 2b. This observation verified for the prediction of most mice but with some exceptions: for instance for feature GLSZM_SmallZone LowGrayLevelEmphasis and the prediction of mouse 8 on the test set, the SHAP values of the images of mouse 8 were not higher than those of the other mice. But mice classification was executed with all features and not only one, so it is not surprising to see such anomalies where one feature alone does not provide a fully accurate prediction of all mice.

**S6 Fig.**
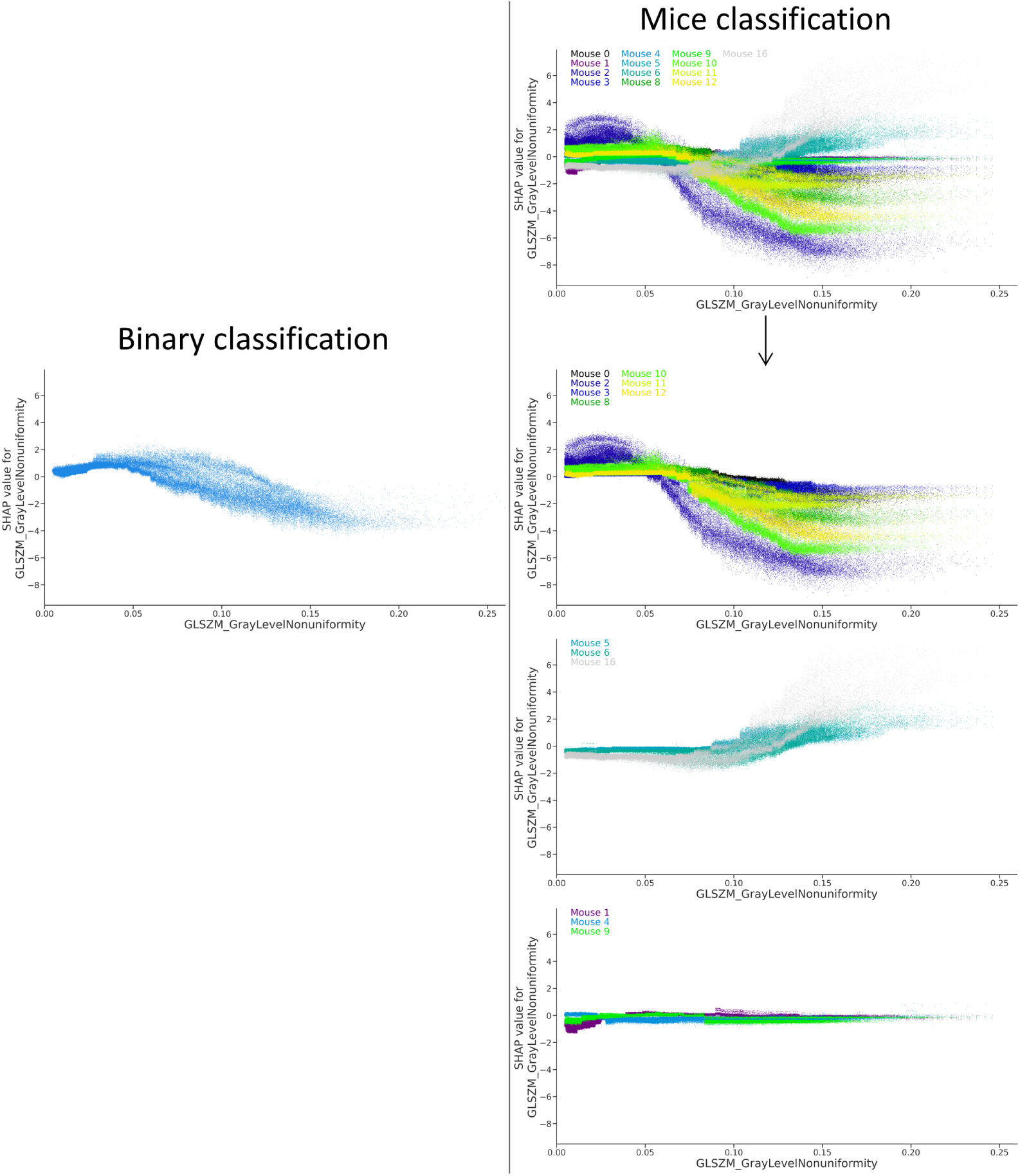
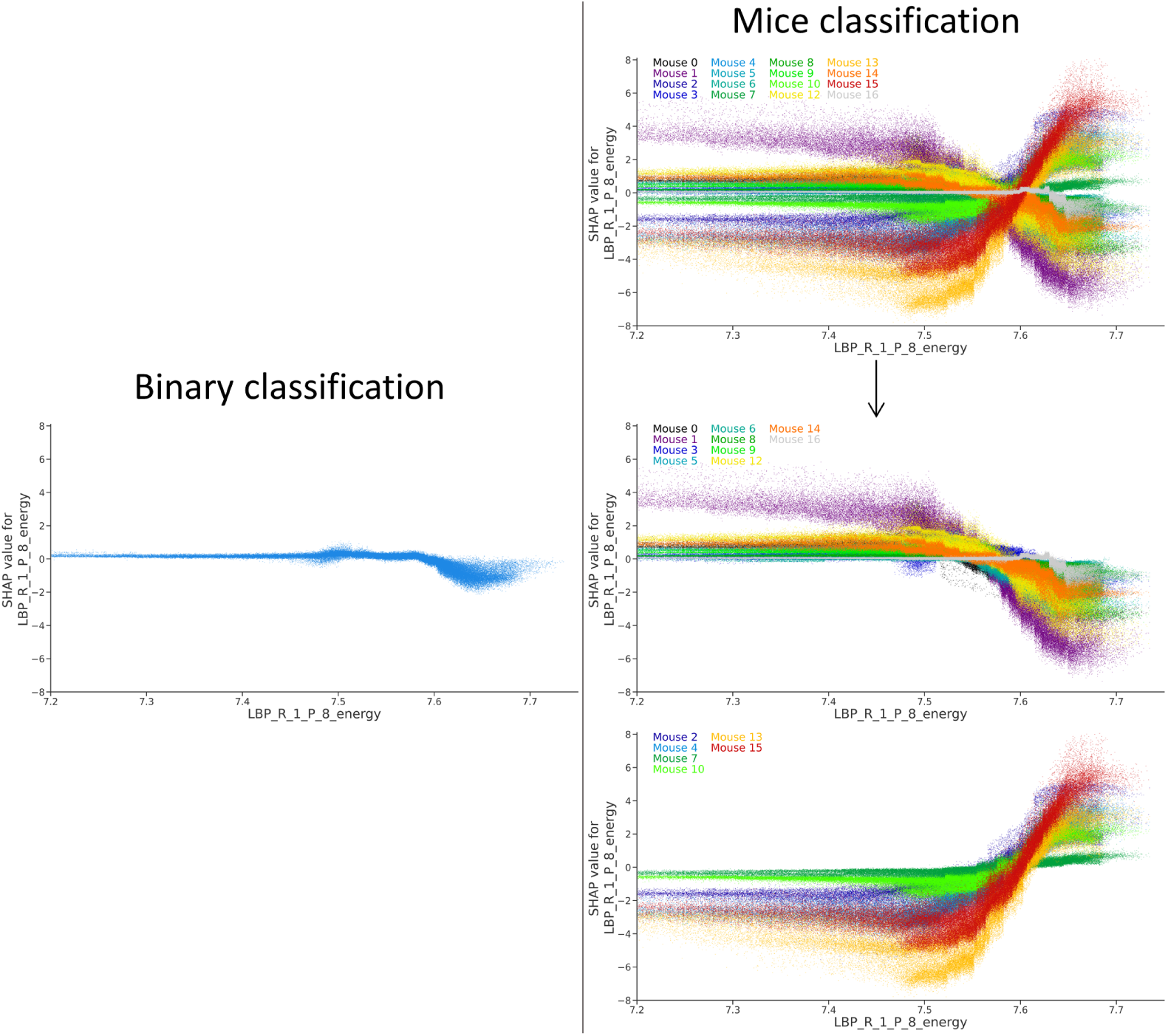
For the most important features, the binary classification dependence plot for the prediction of “regenerating” resembles the mice classification dependence plots of most of the regenerating mice. Dependence plots of features GLSZM_GrayLevelNonuniformity and LBP_R_1_P_8_energy for the XGBoost models trained for binary (left, prediction of “regenerating”) and mice (right, overlaid views) classifications, similarly as in Fig 4. For feature GLSZM_GrayLevelNonuniformity, mice were divided into three groups depending on the shape of their dependence plots. The first group (line two of the right column) was comprised of seven regenerating mice, the second group (line three) of three scarring mice and the third (line four) of two regenerating and one scarring mice. The dependence plot of the XGBoost model trained for binary classification did resemble the superposition of the dependence plots of the first group, which gathered most of the regenerating mice (78%). For feature LBP_R_1_P_8_energy, mice were divided into two groups with similar dependence plots. The first group (line two of the right column) was comprised of six regenerating and four scarring mice and the second (line three) of two regenerating and four scarring mice. The dependence plot of the XGBoost model trained for binary classification did resemble the superposition of the dependence plots of the first group, which gathered most of the regenerating mice (75%).

**S3 Table.**
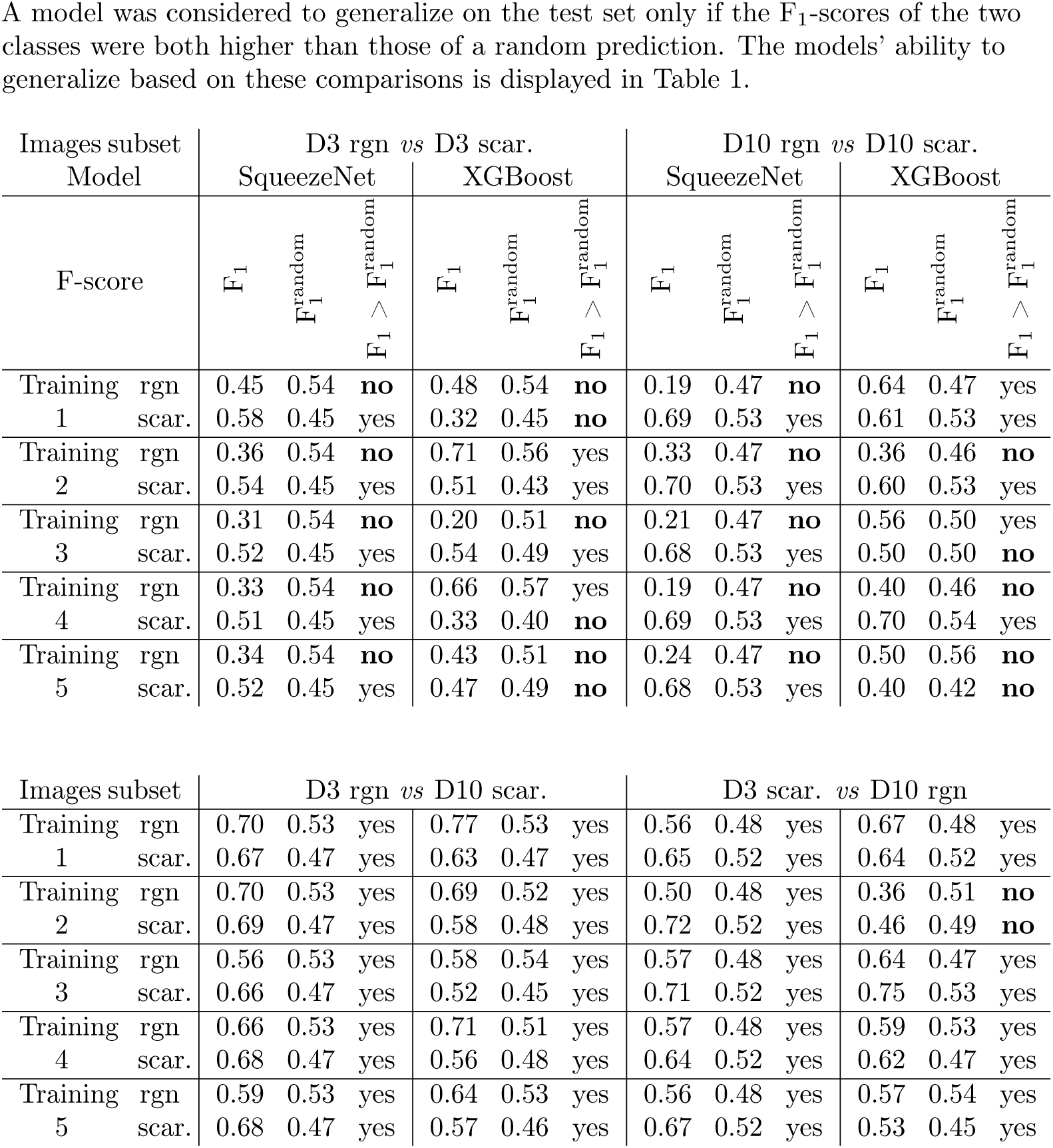
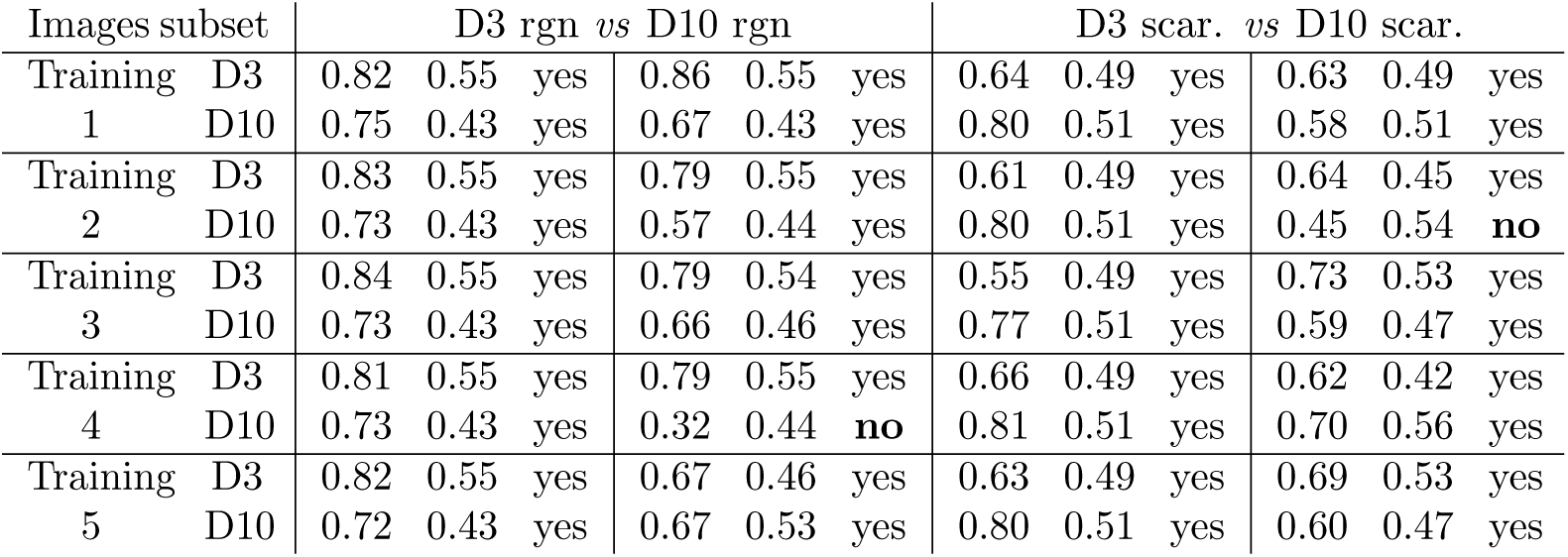
F-scores on the test sets for the deep neural network SqueezeNet and XGBoost on some subsets of images. The two models were trained on the six images combinations of the biological subgroups and then evaluated on the test sets by computing their F_1_-scores for each class. They were compared to the F_1_-scores a random prediction would have produced 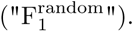 Given two classes *R* and *S* with respectively *r* and *s* images, a random classifier would predict 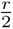 images as *R* and 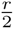 as *S* (resp. 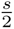 as *S* and 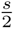 as *R*). The precision of class *R* (resp. *S*) would be 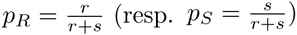 and its recall 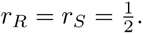 Therefore, the F_1_-score of a random classifier is

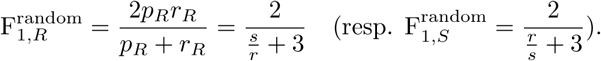 A model was considered to generalize on the test set only if the F_1_-scores of the two classes were both higher than those of a random prediction. The models’ ability to generalize based on these comparisons is displayed in Table 1.

**S7 Fig.**
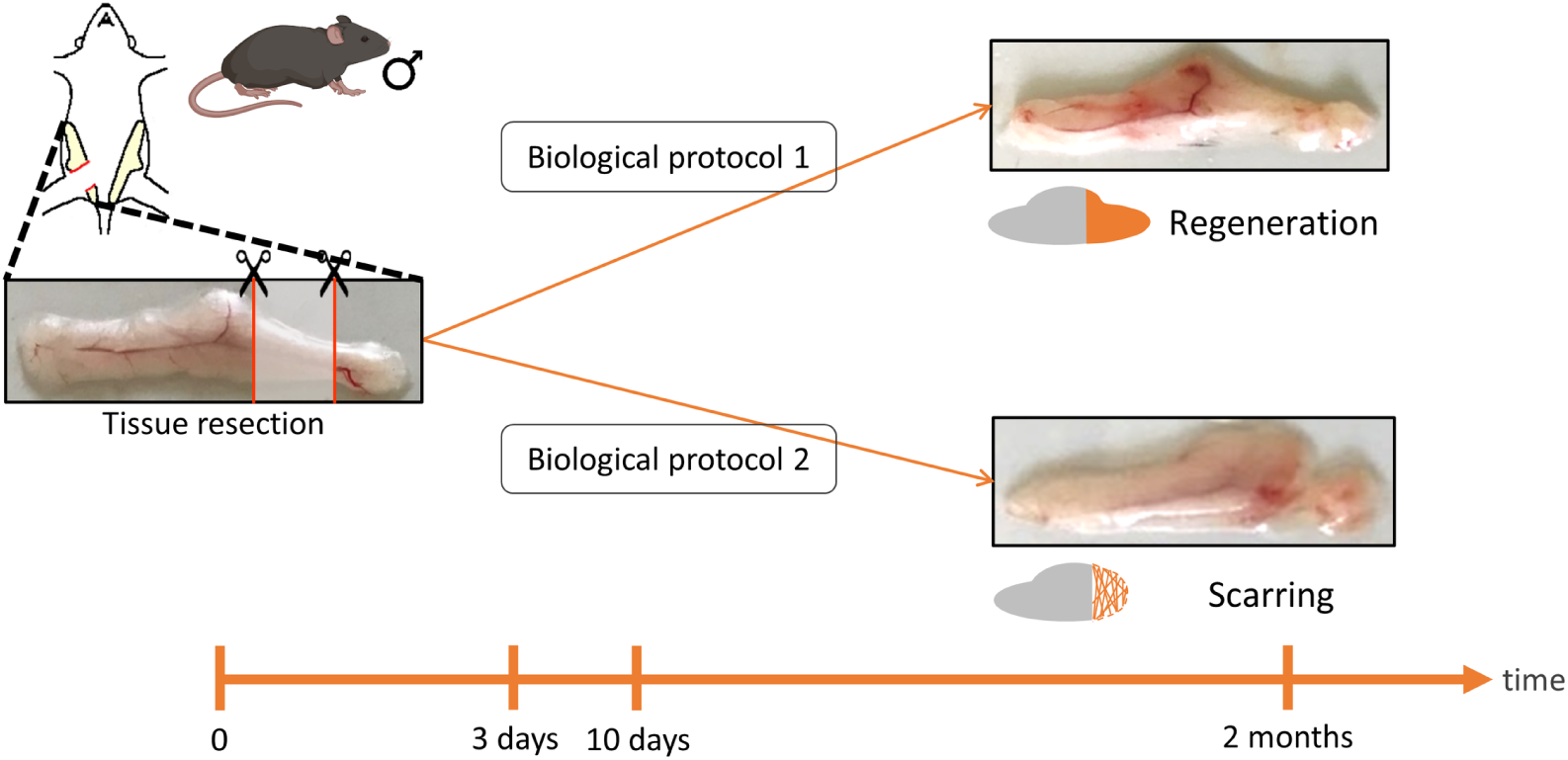
The two biological protocols leading to regenerated or scarred tissues after a two-month wait. Here we use tissues of mice sampled 3 or 10 days after the lesion.

